# Automated MagNet Enrichment Unlocks Deep and Cost-Effective LC-MS Plasma Proteomics

**DOI:** 10.1101/2025.05.06.652407

**Authors:** Erkka Järvinen, Xiaonan Liu, Markku Varjosalo, Salla Keskitalo

## Abstract

Plasma is an ideal material for proteomics due to its diverse protein content reflecting physiological and pathological states, and its compatibility with minimally invasive sampling. Deep proteomic profiling of plasma is hindered by the dominance of high-abundant proteins that mask the detection of low-abundant proteins. To overcome this, we compared five plasma protein enrichment methods, MagNet, ENRICHplus, ENRICHiST, EasySep, and EXONET, against neat plasma using LC-MS proteomics. All five methods substantially increased protein identifications, with MagNet, ENRICHplus, EasySep, and EXONET yielding up to 4200 proteins per sample, over 7-fold more than neat plasma, using a 44-minute gradient on the Evosep One and data-independent acquisition on the timsTOF Pro 2. These methods enriched extracellular vesicle-associated proteins while effectively depleting high-abundant proteins. We further optimized the cost-effective MagNet protocol by increasing the plasma-to-bead ratio and automated the workflow, including Evotip loading, on the Biomek i5 liquid handler. Automated MagNet demonstrated high reproducibility and a remarkably low total cost of just a few dollars per sample. This streamlined enrichment strategy enables scalable, high-throughput LC-MS plasma proteomics, supporting biomarker discovery across large clinical cohorts.

## Introduction

Plasma is a minimally invasive biological material that is typically available in sufficient quantities within clinical studies and large biobank collections. Its wide availability makes plasma an excellent matrix for discovering diagnostic and prognostic disease and treatment markers in untargeted proteomics studies. However, plasma proteomics is complicated by the overwhelming dominance of highly abundant proteins, such as albumin, complement system proteins, and lipoproteins.^1,2^ The concentration range of plasma proteome spans 12 orders of magnitude, with just 22 proteins accounting for 99% of the total protein content.^1,3^

Typical liquid chromatography-mass spectrometry (LC-MS)-based untargeted proteomics of neat plasma identifies around 500 proteins, as most peptides detected by LC-MS are derived from a few hundred highly abundant proteins, hindering the detection of low-abundant proteins.^4–6^ Still, plasma contains large numbers of cellular leakage proteins that are present at significantly lower concentrations than classical plasma proteins.^1–3,7^ Several techniques have been developed to improve protein identifications from neat plasma, including immunodepletion of abundant proteins combined with or without peptide pre-fractionation, and various solvent- or acid-based precipitation workflows.^4,7–9^ However, these methods typically reach no more than ∼1000 protein identifications.^7–9^

Targeted antibody- or aptamer-based affinity proteomics methods, which detect predefined panels of proteins, have gained popularity as alternatives to untargeted LC-MS proteomics.^4^ A recent publication profiled the plasma proteome of over 50,000 adults using an early-generation antibody assay measuring up to 3000 proteins.^10^ The latest version of the aptamer-based assay reportedly covers nearly 10,000 unique protein targets.^11^ A key drawback, however, is that antibodies and aptamers may be affected by epitope variations due to genetic polymorphisms or post-translational modifications, and may suffer from cross-reactivity between targets and thus limiting their specificity compared to LC-MS.^4,12^

Plasma protein enrichment methods have recently expanded protein identifications up to 5000 plasma proteins. These methods include nanoparticles that form protein coronas when mixed with plasma (e.g. Proteograph Assay)^13,14^, hyper-porous magnetic beads with quaternary ammonium groups that bind to phospholipid bilayer-bound particles in (MagNet)^15,16^, and other commercial magnetic particle-based enrichment methods (e.g. ENRICHiST, ENRICHplus, Proteonano).^14,17,18^ With *Lycopersicon esculentum* lectin-mediated plasma enrichment, up to 8500 proteins were reported to be detected from just 25 µl of plasma.^19^ Similarly, extracellular vesicles (EVs) and other extracellular particles carrying tissue-derived proteins have emerged as promising targets to increase plasma proteome depth.^20–23^

Extracting EVs and other extracellular particles from plasma remains a key bottleneck for large scale studies.^24^ Traditional methods such as differential centrifugation or size exclusion chromatography are labor-intensive and difficult to scale for large plasma cohorts. Magnetic bead-based isolation of plasma EVs, by capturing phosphatidylserine or CD9+CD63+CD81 positive EVs, have been proposed as a scalable alternative.^23,25^ These approaches have shown comparable or better performance than differential centrifugation, enabling the identification of more than 4,000 proteins from human plasma.^5,22,23,25,26^ Moreover, magnetic bead-based methods are compatible with automation, making them especially attractive for high-throughput plasma proteomics.

The potential of LC-MS proteomics using data-independent acquisition (DIA) has been demonstrated in recent studies analyzing cohorts ranging from several hundred to thousands of plasma samples, achieving throughput rates of up to 60 samples per day.^8,27–29^ To analyze such large cohorts, automation of sample preparation workflows is essential to reduce manual intervention and improve reproducibility. Fully automated workflows now exist that integrate all key steps of plasma proteomics, including protein extraction, reduction, alkylation, digestion, and peptide loading into disposable trap columns (Evotips), preparing samples directly for LC-MS analysis.^6^ Some systems even support solid-phase peptide purification within the liquid handling platform.^28^

In this study, we systematically evaluated five different plasma enrichment methods, alongside neat plasma, and benchmarked their performance in LC-MS proteomics. We selected the MagNet workflow^15,16^, two commercial enrichment kits (ENRICHiST and ENRICHplus), and two commercial antibody-based EV enrichment beads (EasySep and EXONET). We also assessed the effect of LC gradient length on protein and precursor identifications. Finally, we implemented the MagNet workflow on a liquid handling workstation, streamlining the process from plasma enrichment to Evotip loading for LC-MS-ready samples. This work establishes a scalable, cost-effective framework for high-throughput plasma proteomics, enabling the analysis of thousands of plasma samples in large-scale studies.

## Materials and Methods

### Plasma samples

Blood samples were collected from three female and three male volunteers in EDTA blood collection tubes. Plasma was separated using a single centrifugation at 1500 g for 15 min at room temperature (RT) according to the Early Detection Research Network Standard Operating Procedure for Collection of EDTA Plasma.^30^ The plasma layer was carefully collected from above the buffy coat layer, leaving approximately 1 ml of plasma in each blood collection tube. Plasma samples were gently mixed, aliquoted into 1.5 ml protein low-binding tubes, and transferred to a -80 °C freezer until use. Blood samples were collected as part of the FINPIDD (Hereditary Immune Deficiencies in Finland) study, where volunteers served as healthy adult controls. The study was approved by the Ethics Committee of Helsinki University Hospital. The volunteers provided written informed consent to participate in the study.

Three commercial EDTA plasma samples (PLSSKF2EDT-FSXX, -MSXX, and -XSXX) from a female and two male donors were acquired from Research Donors Ltd (London, UK). Plasmas were thawed either on ice (Commercial 1 and 3) or at RT (Commercial 2), mixed gently, aliquoted into 1.5 ml protein low-binding tubes, and stored in a -80 °C freezer until use.

For each workflow below, plasma aliquots were thawed at RT and briefly vortex-mixed before being subjected to workflows.

### Neat workflow

Neat plasma was prepared according to a previous protocol.^31^ Five microliters of plasma was mixed with 55 µl of 8 M urea (Thermo Fisher Scientific, Waltham, MA, USA) and 10 mM dithiothreitol (DTT, Thermo Fisher Scientific) in 50 mM ammonium bicarbonate. The sample was vigorously vortex-mixed for 1 min, then reduced at 30 °C for 1 hour with 1000 RPM shaking. Proteins were alkylated by adding 2.5 µl of 500 mM iodoacetamide (IAA, Thermo Fisher Scientific) to a final concentration of 20 mM and incubating at RT, protected from light, for 1 hour under 1000 RPM shaking. After alkylation, the sample was diluted with 160 µl of 50 mM Tris-HCl pH 8.0. Fifty microliters of diluted sample were transferred to a new tube containing 60 µl of 50 mM Tris-HCl pH 8.0. Two microliters of 1 µg/µl Trypsin-LysC (Trypsin/Lys-C Mix, Mass Spec Grade, V5071, Promega, Madison, WI, USA) was added to the diluted sample, and digestion was carried overnight at 37 °C under 1000 RPM shaking. After digestion, the sample was acidified with 10 µl of 10% trifluoroacetic acid (TFA, Thermo Fisher Scientific). Finally, samples were diluted to a concentration of 10 ng/µl, based on an estimate of initial plasma protein concentration of 60 µg/µl, by transferring 2 µl of the sample to 130 µl of Solvent A (0.1% formic acid). Subsequently, samples were stored at -80 °C before analysis.

### MagNet workflow

The hyper-porous strong-anion exchange (SAX) magnetic microparticle (MagNet) enrichment method with MagReSyn SAX beads (ReSyn Biosciences, Edenvale, South Africa) has been previously described.^15,16^ The magnetic particles (2.5 µl of 20 mg/ml stock) were aliquoted in a 1.5 ml protein low-binding tube and 200 µl of wash buffer (50 mM BIS-TRIS propane pH 6.3 and 150 mM sodium chloride) was added before a brief shaking at 1000 RPM. The bead washing step was repeated once. Thawed plasma was diluted with an equal amount of binding buffer (100 mM BIS-TRIS propane pH 6.3 and 150 mM sodium chloride). For optimization of the plasma to bead ratio, the volume of beads was kept constant, and either 20, 40, 100, or 200 µl of diluted plasma was applied to the washed beads. The total volume was brought to 200 µl in each case by adding the wash buffer. Plasma and beads were incubated for 30 min at RT with 1000 RPM shaking before the beads were washed three times with 500 µl of the wash buffer, each wash for 5 min at 1000 RPM. Bead-captured proteins were lysed and reduced with 100 µl of 1% sodium dodecyl sulfate (SDS) and 10 mM Tris(2-carboxyethyl)phosphine hydrochloride (TCEP, Thermo Fisher Scientific, Waltham, MA, USA) in 50 mM Tris-HCl pH 8.5 at 37 °C for 1 hour with 1000 RPM shaking. Proteins were further alkylated by adding IAA to a final concentration of 15 mM, followed by incubation at RT for 30 min under 1000 RPM shaking, protected from light. Protein aggregation capture^32^ on the beads was induced by adding acetonitrile to a final concentration of 70%, pipetting the suspension five times up and down, and incubating for 10 min at RT. The beads were washed three times with 95% acetonitrile on a magnetic rack, followed by addition of 200 µl of 50 mM Tris-HCl pH 8.0 containing 750 ng of Trypsin-LysC, and then digested overnight at 37 °C under 1000 RPM shaking. After digestion, TFA was added to a final concentration of 0.5%, and the supernatant containing peptides was transferred to a new tube and frozen at -80 °C prior to peptide quantification and analysis.

### EasySep and EXONET workflows

EasySep Human Pan-Extracellular Vesicle Positive Selection Kit and EXO-NET (EXONET) Pan-Exosome Capture Kit were obtained from STEMCELL Technologies (Vancouver, BC, Canada) and Promega (Madison, WI, USA), respectively. Thawed plasma was centrifuged for 5 min at 10,000 g, and the resulting supernatant was employed in EasySep and EXONET enrichment.

For EasySep, 25 µl of Selection Cocktail (a mixture of anti-human CD9, CD63, and CD81 antibodies) was added to 200 µl of plasma in a 2 ml protein low-binding tube and incubated for 10 min at RT, followed by addition of 50 µl of Releasable RapidSpheres and further incubation for 10 min at RT. Phosphate-buffered saline (PBS) was added to bring the final volume of sample to 2 ml. The sample was pipetted up and down, and beads were separated in a magnetic rack for 5 min. After removing the supernatant, the beads were further washed three times, each with 1700 µl of PBS, by pipet-mixing and separating the beads for 1 min in a magnetic rack.

For EXONET, 15 µl of Pan-Exosome Capture beads (immunoaffinity magnetic beads with proprietary antibodies) was added to 200 µl of plasma in a 2 ml protein low-binding tube, and the sample was incubated for 15 min at RT. The tube was placed in a magnetic rack for 5 min, supernatant was removed, and beads were washed three times with 1000 µl of PBS by incubating the tube for 5 min in a magnetic rack between washes.

Captured vesicles and proteins onto beads were lysed with 100 µl of 5% SDS in 50 mM TRIS pH 8.5 for 15 min at RT under shaking at 1200 RPM. Beads were separated in a magnetic rack for 5 min, and the supernatant containing lysed proteins was transferred to a new tube. Proteins were reduced by adding TCEP to a final concentration of 5 mM, incubating for 15 min at 55 °C under 600 RPM shaking, followed by alkylation with 20 mM IAA for 10 min at RT, protected from light and under 600 RPM shaking.

Proteins were purified and digested with S-Trap mini spin columns according to the manufacturer’s protocol (ProtiFi, Fairport, NY, USA). The sample was acidified to a pH below 1 with phosphoric acid at a final concentration of 1.1%. Binding buffer (100 mM TRIS-HCl pH 7.55 in 90% methanol) was added to the tube to be six times the sample volume, and the sample was loaded onto the S-Trap mini spin columns by several centrifugations at 4000 g for 30 seconds. The column was washed four times with 400 µl of binding buffer by centrifuging for 30 seconds at 4000 g, followed by a final centrifugation of empty column for 1 min at 4000 g. Proteins were digested by applying 125 µl of 10 µg Trypsin-LysC in 50 mM ammonium bicarbonate to the column and incubating overnight in a heat cabinet set to 37 °C. After digestion, peptides were eluted by applying 80 µl of 50 mM ammonium bicarbonate, 80 µl of 0.2% formic acid, and 80 µl of 50% acetonitrile into the column and centrifuging between the solvents for 1 min at 4000 g. The resulting flowthroughs were combined and dried in a vacuum centrifuge, reconstituted in 100 µl of Solvent A, and frozen at -80 °C prior to peptide quantification and analysis.

### ENRICH-iST and ENRICHplus workflows

ENRICH-iST and ENRICHplus kits were obtained from PreOmics (Planegg/Martinsried, Germany), and plasma enrichment was conducted according to the manufacturer’s protocols. Briefly, 25 µl of EN-BEADS or ENplus-BEADS were washed twice with wash solvent. Then, 80 µl of EN-BIND or 50 µl of ENplus-BIND was added to the beads, followed by adding 20 µl of plasma (ENRICH-iST) or 50 µl of plasma (ENRICHplus). After 30 min incubation at 30 °C, 100 µl of EN-BIND or ENplus-BIND was added to the sample, followed by two washes with 100 µl of buffer. Samples were lysed, reduced, and alkylated with either 50 µl of LYSE-BCT at 95 °C for 10 min (ENRICH-iST) or 40 µl of LYSE-BIND at 60 °C for 10 min (ENRICHplus). After cooling the samples to RT, 50 µl of DIGEST (ENRICH-iST) or 10 µl of DIGEST (ENRICHplus) was added to the samples, followed by 1 hour incubation at 37 °C. STOP solution (100 µl for ENRICH-iST or 60 µl for ENRICHplus) was added to the samples, and the samples and beads were transferred to purification cartridges. After washing the cartridges with two different wash solutions, peptides were eluted in a total volume of 200 µl, dried in a vacuum centrifuge, reconstituted with 15 µl of LC-LOAD, and frozen at -80 °C prior to peptide quantification and analysis.

### Automated MagNet workflow

The automated MagNet workflow and Evotip loading were implemented to Biomek i5 Automated Liquid Handler from Beckman Coulter (Brea, CA, USA) equipped with a 1200 µl 96-channel head, a Magnum FLX magnet plate from Alpaqua Engineering (Beverly, MA, USA), and a BioShake 3000-T elm heater shaker mounted with a microplate adapter (Microplate adapter - 96 well Bio-Rad PCR plate, part number 2016-1042), both from QINSTRUMENTS (Jena, Germany).

MagReSyn SAX beads were washed with the wash buffer before diluting them to 10-fold higher volume with the wash buffer and aliquoting manually 10 µl of diluted beads, corresponding to 0.5 µl of original bead stock, per well in a 96-well plate (twin.tec PCR Plates LoBind - PCR Plates, product number 0030129512, from Eppendorf (Hamburg, Germany)). Plasma samples, 20 µl, were pre-diluted with 20 µl of the binding buffer, and 40 µl of diluted samples were transferred to the well plate bringing the total well volumes to 50 µl. Alongside, a plate with lysis/reduction/alkylation solution and a plate with digestion solution were prepared. The lysis solution contained 1% SDS, 5 mM TCEP and 10 mM chloroacetamide (Sigma-Aldrich, St. Louis, MO, USA) in 50 mM Tris-HCl pH 8.5, and 20 µl of this solution was aliquoted to each well of a 96-well PCR plate (product number AB0800, Thermo Fisher Scientific). The digestion solution contained 90 ng of Trypsin/Lys-C Mix in 50 µl of 50 mM Tris-HCl pH 8.0, and 55 µl of this solution was aliquoted to each well of LoBind 96-well PCR plate. The plates and three 90 ml reservoirs (product number 242811 from Thermo Fisher Scientific) filled with the wash buffer, isopropanol or Solvent A were transferred to the automated liquid handler and the workflow was performed as reported in Supplementary Table S1.

For the Evotip loading (EV1183 – Evotip Loading Kit for Biomek i5/i7 1200 μL 96-Channel head from Evosep), 15 µl of Solvent A was transferred into tips followed by conditioning the tips in isopropanol for 30 s before a positive pressure push for 4 s to push Solvent A through the resin. Twenty microliter of the diluted peptide samples from the MagNet workflow were transferred to Evotips followed by an air gap and 150 µl of Solvent A. The sample and air gap were pushed through the Evotip resin by positive pressure for 100 s.

The automated MagNet workflow was run in two batches of 96-well plates. Both batches included six quality control (QC) samples, which consisted of 50 ng Pierce HeLa Protein Digest Standard (Thermo Fisher Scientific) loaded manually to Evotips. The QC samples were the first and last LC-MS run, and the remaining four QC samples divided equally during the batch.

### Peptide quantification

The Pierce Quantitative Colorimetric Peptide Assay (Thermo Fisher Scientific) was employed for the peptide quantification in a 96-well plate format according to the manufacturer’s protocol. Each measurement included a peptide standard curve spanning a range of 15.6 ng/µl to 1000 ng/µl prepared from Peptide Digest Assay Standard. Twenty microliters of standards, samples, and blanks (same composition of solvents as in samples) were pipetted into a well plate, followed by 180 µl of Working Reagent. The plate was briefly mixed for 30 seconds at 600 RPM and then incubated for 20 min at 37 °C in a heat cabinet. The absorbance of samples was measured at 480 nm with a CLARIOstar microplate reader (BMG LABTECH, Ortenberg, Germany).

### LC-MS/MS

Peptide samples were loaded into Evotips according to the manufacturer’s protocol by conditioning tips with 20 µl of 0.1% formic acid in acetonitrile followed by soaking them in 1-propanol. The tips were equilibrating with 20 µl of solvent A before loading 10 to 30 µl of samples. The tips were further washed with 20 µl of solvent A before briefly centrifuging them with 100 µl of solvent A and leaving the rest of the solvent A in the tips. Sample loads were 100, 200, and 300 ng of peptides for 100 samples-per-day (SPD), 60 SPD, and 30 SPD methods, respectively. For neat plasma analysis, sample loads were 10, 20, and 30 µl for the 100 SPD, 60 SPD and 30 SPD methods, respectively. The Evosep One liquid chromatography (LC) system^33^ connected to a trapped ion mobility quadrupole time-of-flight MS (Bruker timsTOF Pro 2, Bruker Daltonics, Billerica, MA, US) equipped with CaptiveSpray nano-electrospray ion source (Bruker Daltonics) was employed for the LC-MS analysis. A 15 cm × 150 μm column with 1.5 μm C18 beads (EV1137, Evosep) was employed for the 30 SPD (44 min gradient) method. An 8 cm × 150 μm column with 1.5 μm C18 beads (EV1109, Evosep) was employed for the 60 SPD (21 min gradient) and 100 SPD (11.5 min gradient) methods. The column oven was set to a temperature of 40 °C for all methods. Mobile phases A and B were 0.1% formic acid in water and 0.1% formic acid in acetonitrile.

For the MS data acquisition, DIA with parallel accumulation-serial fragmentation (dia-PASEF)^34^ was used with default source parameters and optimized isolation windows (Supplementary Tables S2-S6).

### MS data analysis

Raw dia-PASEF data were processed with DIA-NN version 1.9.2.^35,36^ A spectral library was generated from a human FASTA file (downloaded on 08/11/2024 from UniProtKB/Swiss-Prot database, containing 20,404 proteins) using the following precursor ion generation parameters: Trypsin/P (protease), 1 missed cleavage, cysteine carbamidomethylation enabled as a fixed modification, a maximum of 3 variable modifications with N-terminal methionine excision, methionine oxidation, and N-terminal acetylation enabled, peptide length range from 7 to 30, precursor charge from 2 to 4, 294-1325 m/z precursor range, and 100-1700 m/z fragment ion range. For the raw file searches, heuristic protein inference was enabled, and protein inference was set to protein names from FASTA. Mass accuracies were set to 15 ppm and the scan window to 7 (100 SPD), 8 (30 SPD), or 10 (60 SPD). Unrelated runs option was not selected, while peptidoform scoring and match-between-runs were selected, and IDs, RT, and IM profiling was chosen for the library generation. Machine learning, quantification strategy, and cross-run normalization were set to single-pass NNs, QuantUMS, and RT-dependent. The false discovery rate for the output filtering was set to 1%.

### Data analysis

Protein identification results from the 30 SPD method runs were used for all analyses except Figures 1 and S2 that report results for the gradient comparisons, and Figures 4B, 4D and S12 that report results from 100 SPD runs. Protein groups with multiple entities and keratins (common contaminants) were excluded from all analyses, except Figures 1, 2A, 4B, 4D, S2 and S12. Commercial plasma samples were included only in the analyses presented in Figures 2 and S2. For hierarchical clustering, the Euclidean distance measure and Ward’s clustering method were applied. The PANTHER database^37,38^ was employed for human Gene Ontology (GO), pathway, and protein class ontology enrichment analyses with 5% false discovery rate filtering, further refined by selecting terms with at least 2-fold enrichment and containing at least 10 proteins per term. The subcellular location of proteins, gene-level RNA expression data, single cell-type expression data, and the human secretome annotations were downloaded from the Human Protein Atlas (HPA, version 24.0, proteinatlas.org). All analyses were done in the R programming language environment (R version 4.4.2).

**Figure 1.**
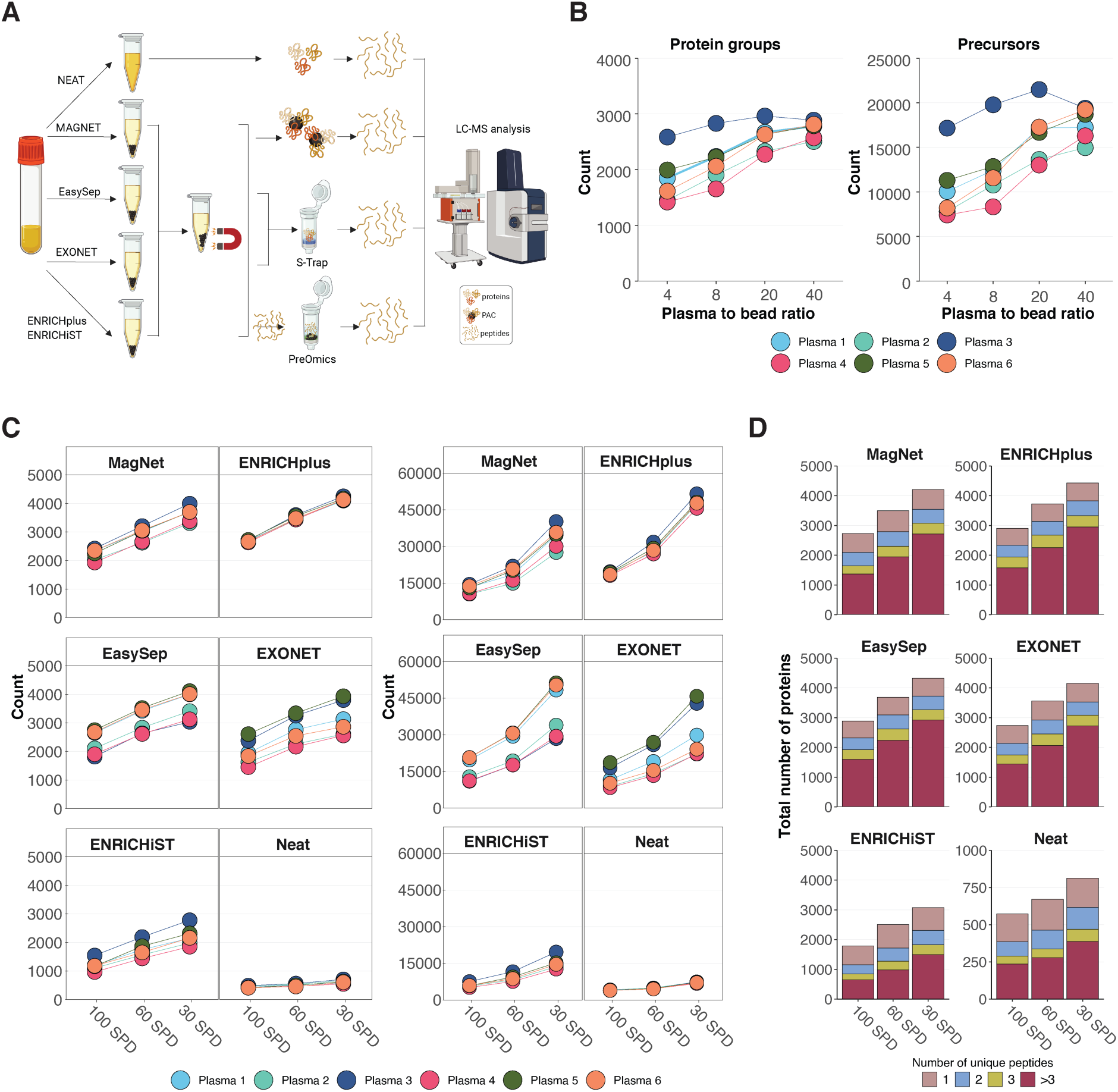
Plasma enrichment methods increase the protein identifications in comparison to neat plasma. **(A)** Overview of the methods used in this study, where five different plasma enrichment methods were compared with neat plasma. **(B)** Optimization of the plasma-to-bead ratio for MagNet using the 60 SPD chromatographic gradient method. The 40:1 plasma-to-bead volume ratio was selected for MagNet samples presented in panels C and D. **(C)** Five different enrichment methods and the neat plasma workflow were applied to six different plasma samples, which were analyzed using three different chromatographic gradients. **(D)** Total number of proteins identified per plasma method and chromatographic gradient, along with the unique peptides identified for each protein.

**Figure 2.**
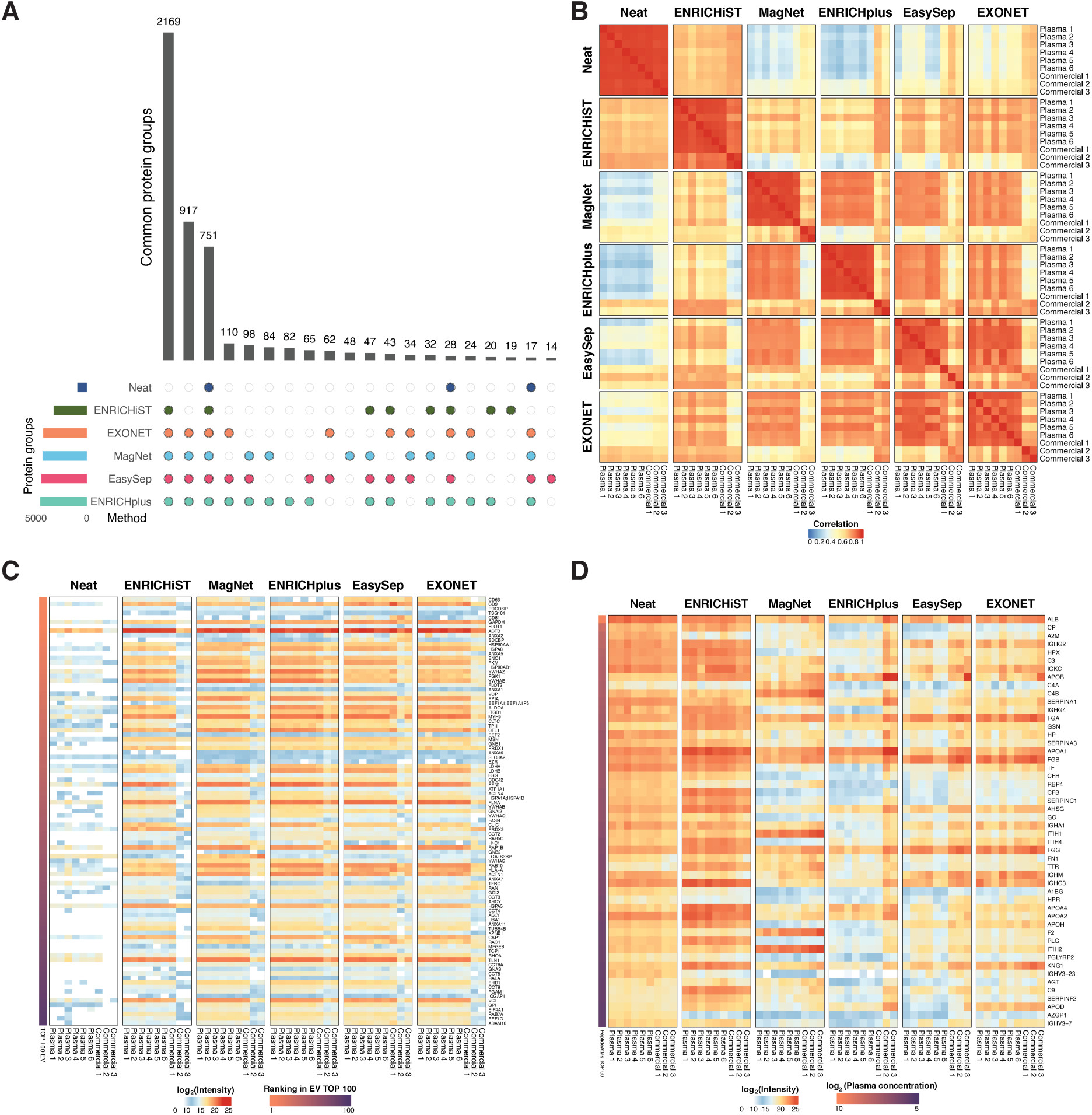
Enrichment methods yield overlapping protein profiles, enriching EV proteins while depleting abundant plasma proteins, with higher correlation within than between methods. **(A)** Overlap of total protein group identifications across different methods. The number of overlapping groups are restricted to total number of 13 proteins. **(B)** Pearson correlation matrix of protein intensities across all plasma samples. **(C)** Log_2_ intensity of the top 100 EV proteins from Vesiclepedia^40^ across all samples. A2M, ALB, C3, FN1 and GSN were excluded from the analysis, as these proteins are also among the top 100 plasma proteins in PeptideAtlas^41^. **(D)** Log_2_ intensity of the top 50 plasma proteins from PeptideAtlas^41^ across all samples. IGHG1, IGHA2, IGHV3-48 and PRSS1 were excluded from the analysis, as these proteins were either not detected or were present in less than 20% of samples. The commercial plasma samples were included only in this figure.

## Results

### Comparison of plasma enrichment methods with neat plasma

We selected five different plasma enrichment methods and compared them with neat plasma, which was processed without any depletion or enrichment (Figure 1A). The selected methods included two commercial enrichment kits, ENRICHplus and ENRICHiST, for which the manufacturer does not specify a precise biochemical mechanism, but claims that low-abundant proteins are enriched onto paramagnetic beads. The other two commercial methods, EXONET and EasySep, rely on antibody-mediated capture of EVs from plasma via proteins on the vesicle surface. For EasySep, the manufacturer specifies that the beads are conjugated with antibodies against CD9, CD61, and CD81. The last method, MagNet, is a recently developed approach that utilizes hyper-porous strong anion-exchange magnetic microparticles, which are reported to enrich phospholipid bilayer-bound particles (e.g. EVs) from plasma.^15,16^

First, we optimized the plasma-to-bead ratio for MagNet. A previous study used a 4:1 plasma-to-bead ratio, however, our results showed that protein and precursor identifications increased up to a 40:1 ratio, where the increase began to plateau (Figure 1B). Therefore, we selected the 40:1 ratio for all subsequent MagNet experiments.

All five enrichment methods increased protein identifications compared with neat plasma when tested across six individual plasma samples (Figure 1C). Additionally, we evaluated three chromatographic gradients and observed a linear increase in protein identifications with longer gradients. Notably, precursor numbers increased even more substantially with the longest gradient 30 SPD (44 min). Longer chromatographic gradients also allowed for higher peptide loading. As shown in Figure S1, the total ion chromatogram (TIC) signal intensity remained consistent across peptide loads of 100, 200, and 300 ng for the 100, 60, and 30 SPD gradients, indicating that the dynamic range of system was not compromised by increased sample input. It is worth noting that our MS instrumentation (timsTOF Pro 2) saturates at peptide loads exceeding 300 ng.^39^

The 30 SPD gradient also employes a 15 cm LC column, whereas the 60 and SPD gradients use an 8 cm LC column. The longer column likely contributes to the substantial gain in precursor identifications observed with the 30 SPD gradient compared to the 60 and 100 SPD methods, though this did not translate into a proportionally higher number of protein identifications (Figure 1C). Additionally, all three gradients yielded sufficient precursor coverage to allow identification of more than one unique peptide per protein in most cases when using enrichment methods (Figure 1D). For example, the top-performing enrichment methods (MagNet, ENRICHplus, EasySep, and EXONET) yielded more than one unique peptide for over 80% of identified proteins (Figure 1D).

In addition to the six plasma samples collected from our healthy control cohort, we included three commercial plasma samples in the analysis. Comparing the enrichment methods between commercial and other plasma samples revealed that two of the three commercial samples had at least two-fold fewer protein and precursor identifications (Figure S2). Similar trends were observed in the correlation analysis (Figure 2B) and the analysis of the top 100 EV proteins (Figure 2C), where Commercial 2 and 3 plasma samples appeared as clear outliers. Although the commercial plasma samples were derived from EDTA-treated blood similarly to other plasma samples, we lack exact details on their preparation and thus cannot speculate on the reasons for these discrepancies. Therefore, we excluded the three commercial samples from all subsequent analyses.

### Correlation between methods, enrichment of EV proteins and depletion of high-abundant plasma proteins

Most proteins identified in this study, 3837 out of 4865 protein groups, were detected using the four most effective enrichment methods: MagNet, ENRICHplus, EasySep, and EXONET (Figure 2A). ENRICHiST was the least effective, identifying only 3232 of the 4865 total protein groups. Method-specific protein identifications were minimal, and for example 82 protein groups were unique to ENRICHplus, while 48 protein groups were specific to MagNet, highlighting the high similarity between the evaluated enrichment methods. Protein intensity correlations revealed strong intra-method correlations (Figure 2B). Notably, ENRICHiST samples correlated closely with neat plasma, suggesting limited enrichment efficacy. In contrast, the other four enrichment methods showed moderate correlation among themselves but substantially lower correlation with neat plasma.

To assess EV protein enrichment, we analyzed the intensity of the top 100 EV proteins from Vesiclepedia^40^ across all samples (Figure 2C). All five enrichment methods similarly enriched EV proteins, which were absent or detected at low intensity in neat plasma samples. Next, we examined the 50 most abundant plasma proteins, based on estimated concentrations from PeptideAtlas^41^ (Figure 2D). MagNet, ENRICHplus, EasySep, and EXONET effectively depleted most high-abundant plasma proteins compared to neat plasma, whereas ENRICHiST showed limited depletion and even enriched some of these proteins, aligning with its weaker correlation to other enrichment methods (Figure 2B).

Most proteins identified with enrichment methods overlapped with the current Human Plasma and Human Plasma EV builds in PeptideAtlas (Figure S3). However, 600–800 proteins detected with the four most effective enrichment methods had not been previously reported in these datasets. Additionally, the enrichment methods enabled detection of proteins across a wide dynamic range (∼10⁹-fold) of reported human plasma protein concentrations (Figure S3).

### Comparison of enrichment methods to previous plasma studies

We compared our MagNet results to a previous MagNet study^15^ and observed a high degree of overlap in protein identifications, along with a strong correlation between protein intensities across both datasets (Figures S4A and S4B). Similarly, comparison of our ENRICHiST data with a prior study using the same method^14^ showed that over 50% of proteins overlapped, but we identified 1417 additional proteins not previously reported with this method (Figure S4C). Finally, MagNet, ENRICHplus, EXONET, and EasySep yielded a similar total number of protein identifications compared to Seer Proteograph XT, a widely adopted commercial enrichment assay.^14^ However, nearly half of the identified proteins were unique to either to Seer Proteograph XT or the enrichment methods used in this study (Figure S4D), highlighting method-specific biases in protein enrichment.

Comparisons with previous studies analyzing plasma EV proteins demonstrated a high overlap between our results and previous EV proteomics datasets (Figures S5A-C). Furthermore, protein intensity comparisons between our enriched samples and two earlier studies^5,26^, that isolated EVs from platelet-poor plasma via centrifugation, revealed good correlation between (Figures S5D and S5E). However, the correlation was markedly weaker when comparing our data to phosphatidylserine-positive EVs isolated from filtered plasma^23^ (Figure S5F), suggesting that different EV isolation methods capture distinct extracellular particle populations. Finally, we confirmed that a major proportion of proteins detected using the enrichment methods are included in the commercial Olink HT and SomaScan 11K targeted protein assays, although approximately a quarter of the proteins identified using the four most effective enrichment methods in this study are not covered by these targeted platforms (Figure S6).

### Clustering and enrichment analysis of enriched proteins

We filtered all identified proteins to include only those detected in at least five samples, resulting in a subset of 4623 proteins, which we clustered based on their intensity across all samples. This analysis grouped the proteins into four main clusters (Figure 3A), corresponding to their detection patterns across the enrichment methods. Cluster 1 contained proteins consistently detected by all methods, including most proteins present in neat plasma. Cluster 2 included highly abundant proteins enriched by all five enrichment methods. Cluster 3 consisted of proteins of moderate intensity, primarily enriched by MagNet, ENRICHplus, EasySep, and EXONET. Cluster 4 contained mostly low-abundant proteins that appeared inconsistently across enrichment methods.

**Figure 3.**
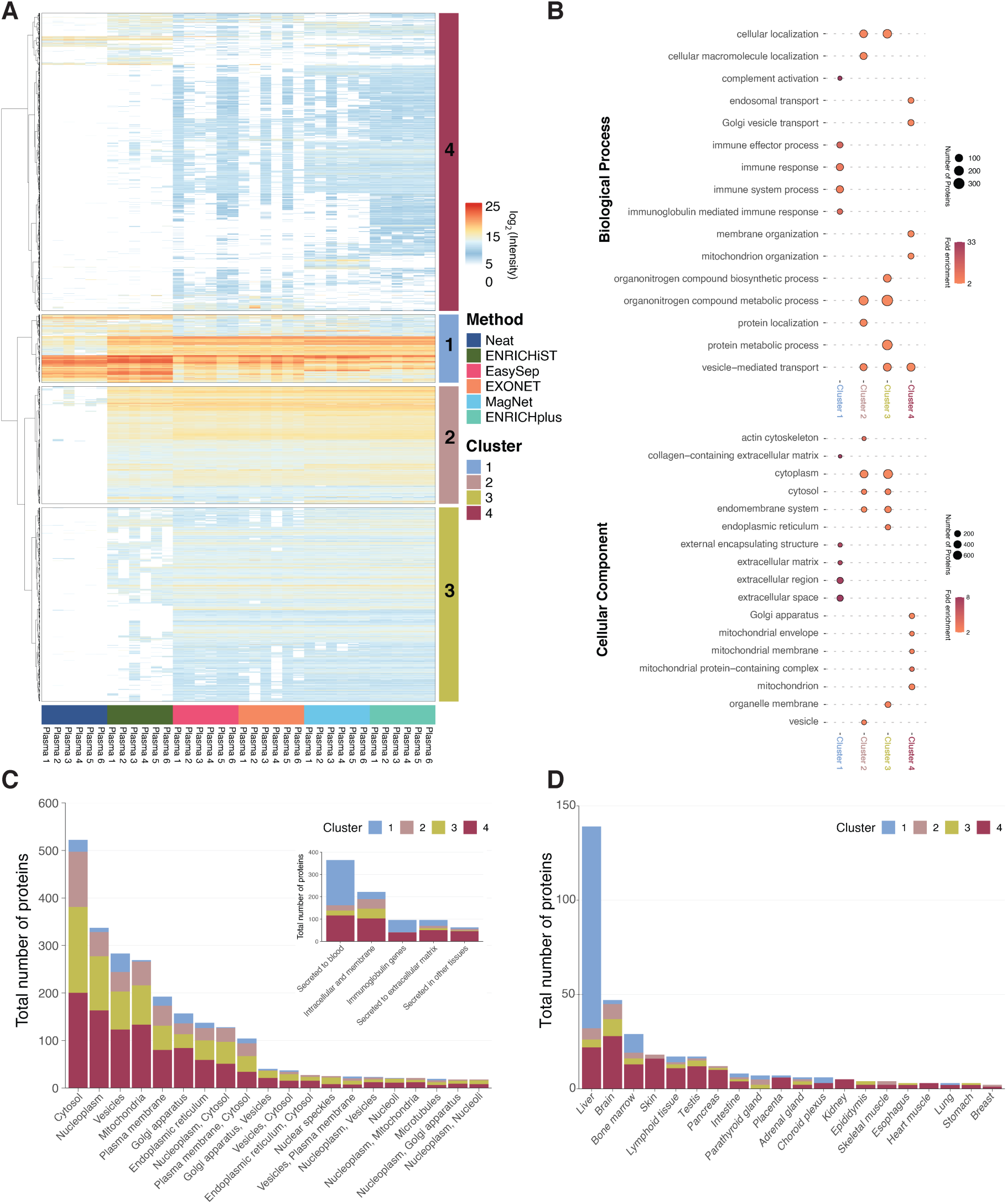
Enriched and neat plasma proteins cluster into four main groups based on protein intensities. Cluster 1 contains proteins found in neat plasma and shared across all methods, while Cluster 2 consists of proteins identified in all five enrichment methods. Cluster 3 includes proteins with medium intensity, mostly detected in all methods except ENRICHiST and neat. Cluster 4 consists primarily of low-intensity proteins. Each cluster is enriched with distinct Gene Ontology terms, cellular localizations and tissue associations. **(A)** Hierarchical clustering of proteins detected in at least 10% of all samples, based on log₂ intensity. **(B)** Enrichment analysis of proteins in each cluster with the PANTHER GO-Slim databases^37,38^ for cellular components and biological processes. Results were first filtered to retain terms with over two-fold enrichment and a minimum of ten proteins per term. The top five terms with the lowest FDR values were selected for each cluster. **(C)** Subcellular location of proteins in each cluster based on The Human Protein Atlas (HPA) annotations.^42^ Only the top 20 locations are shown. The inset highlights the top five secretome^2^ annotations among all identified proteins. **(D)** Number of proteins annotated as tissue-enriched in the HPA transcriptomics dataset^43^. The 20 tissues with the highest total count of tissue-enriched proteins are shown.

We performed GO analysis for each cluster (Figures 3B and S7), along with pathway and protein class enrichment analyses (Figure S7). Cluster 1, mainly composed of proteins found in neat plasma, showed enrichment for biological processes related to blood coagulation, hemostasis, immune system function, and immunoglobulins (Figures 3B, S7, and S8). These proteins primarily localize to the extracellular region, space and matrix (Figure 3B), consistent with their role as circulating plasma proteins and the main component of neat plasma proteins. Clusters 2 and 3 were enriched for for cytosolic, cytoplasmic, and vesicle-associated proteins, particularly those involved in metabolic and signaling pathways (Figures 3B, S7, and S8). In contrast, Cluster 4 had a distinct enrichment profile, featuring proteins associated with mitochondrial, Golgi, and endosomal transport functions (Figure 3B). The differences between four clusters highlight the biological diversity of proteins captured by plasma enrichment methods.

To further characterize the protein profiles, we analyzed the subcellular localization^42^, predicted secretion^2^, and tissue-specific transcript expression^43^ of each cluster (Figures 3C and 3D). Most detected proteins localize to the cytosol and nucleoplasm, though a substantial number of membrane proteins were also identified (Figure 3C). All clusters displayed similar distributions across subcellular compartments, though Clusters 3 and 4 contained the highest number of proteins in each category. Secretome analysis revealed that blood proteins and immunoglobulins were mainly present in Cluster 1, while intracellular and membrane proteins were predominantly enriched in Cluster 4 (Figure 3C, inset).

We assessed tissue-specific expression, which showed that most tissue-enriched proteins originated from the liver and bone marrow and were primarily part of Cluster 1 (Figure 3D). Other proteins annotated as tissue-enriched in the HPA transcriptomics dataset were mainly in Cluster 4 that contained low-intensity plasma proteins (Figure 3A). Analysis of cell-type-specific proteins using the HPA single-cell transcriptomics dataset confirmed that plasma cells and hepatocytes, major sources of blood proteins, were the most enriched (Figure S9).

To assess tissue specificity at both transcript and protein levels, we analyzed tissue-enriched transcripts from the HPA RNA dataset^43^ (Figure S10A), and tissue-enriched proteins from ProteomicsDB^44^ and The Human Proteome Map^45^ datasets (Figures S10B and S11A). Consistent with the findings in Figures 3D and S9, liver, bone marrow, skin, and testis emerged as the most distinct tissues in the HPA RNA dataset (Figure S10A). In contrast, proteomics datasets (Figures S10B and S11A) did not highlight the liver as the most prominent tissue, likely because secreted proteins are less enriched at the protein level. Instead, blood immune cells and platelets showed the strongest enrichment for proteins detected in our dataset (Figure S11A).

Finally, we assessed the correlation between proteins enriched in our study and those detected in The Human Proteome Map dataset^45^, focusing on blood immune cells and the liver (Figure S11B). Among the over 2000 shared proteins, those enriched by MagNet, ENRICHplus, EasySep, and EXONET showed strong correlation (R^2^ = 0.45-0.49) with the platelet proteome, while correlations with other blood cell types and liver were clearly lower (Figure S11B).

### Automated plasma enrichment workflow

We developed an automated plasma enrichment method for the Biomek i5 liquid handler based on the MagNet workflow, fully integrating it with a novel, automated Evotip loading method. We selected the MagNet workflow for automation due to its significantly lower cost - over an order of magnitude cheaper than the other enrichment methods (Table S7). Furthermore, it is only workflow that does not require cartridge-based purification. The automated Evotip loading method performed comparably to manual loading (Figures S12A and S12B).

To evaluate the performance of automated MagNet workflow, we processed two 96-well plates, each containing 15 replicates of six plasma samples, and 6 replicates of blank samples without plasma (Figure 4A). In addition, we included six QC samples for both LC-MS runs, prepared by manually loading 50 ng of HeLa protein digest standard into Evotips. The QC samples demonstrated consistent performance across both batches (Figure S12B). The coefficient of variation (CV) of protein quantification across the full MagNet workflow, from plasma sample to quantified protein values, was well below 20% both within and between plates (Figure 4B). The CVs of precursor quantification were higher but did not exceed 25% in any case (Figure S12C). Although some proteins were identified in a few blank samples, their total precursor counts never exceeded 0.5% of the average sample count.

**Figure 4.**
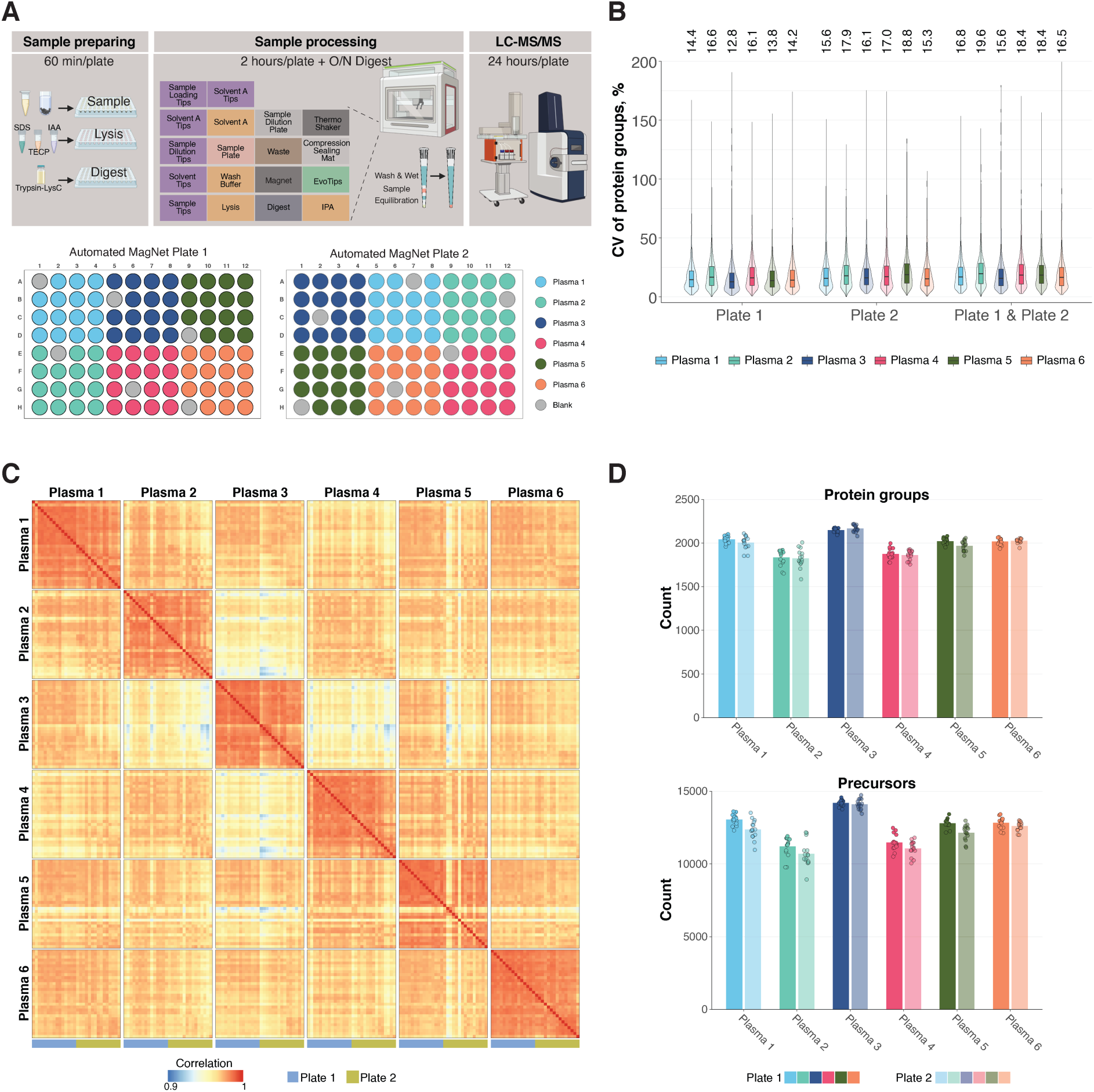
Automated MagNet workflow with combined Evotip loading using a Biomek i5 liquid handler shows high reproducibility. **(A)** Overview of the automated MagNet and Evotip workflows, and plate layouts for two MagNet runs. The 100 SPD method was used for LC-MS analysis of all samples. **(B)** Coefficient of variation (CV, %) of protein groups within each plasma sample across plate 1 and plate 2 (n = 15 per plate; n = 30 total). The exact values of median CVs are shown above the plot **(C)** Pearson correlation of protein intensities between plasma samples across both plates. The scale bar is from 0.9 to 1. **(D)** Number of identified protein groups and precursors per plasma sample on each plate (n = 15 per plate; n = 30 total).

The correlation of protein quantification within the same plasma samples was mostly between 0.98-0.99, and no differences were observed between the plates, indicating the absence of batch effects or well position biases (Figure 4C). A few samples, particularly in Plasma 2 and Plasma 5, showed slightly lower correlation of 0.95-0.97 to the rest of samples within the same plasma, but there was no clear explanation for these slightly lower correlations, such as edge effects or reduced MS1 signal (Figure S12D).

MS1 signal intensity remained largely stable, with the exception of the first ten runs of Plate 2, which showed slightly elevated intensity compared to Plate 1 and the remaining samples (Figure S12D). However, this variation did not affect sample correlation or protein/precursor identifications (Figures 4C and 4D). The elevated MS1 signal was likely due to a plate edge effect, as wells in row A of Plate 2 appeared to contain a reduced volume, despite proper sealing with foil.

The total number of protein groups and precursors identified was highly consistent across replicates (Figure 4D). While protein group identification was approximately 200 proteins lower than in the manual workflow (Figure 1C), it is important to note that results in Figure 1C were part of a same DIA-NN search, where the complementary nature of different enrichment methods and gradients likely boosted identifications via match-between-runs. Finally, we evaluated the digestion efficiency of the automated MagNet workflow, as this step initially required multiple rounds of optimization. Digestion efficiency was consistent across all samples and both plates, with a mean value of 16% and a range from 14% to 18% (Figure S12E). For comparison, the manual MagNet workflow yielded an average digestion efficiency of 14% (n = 9).

## Discussion

The main finding of this study is that the four plasma enrichment methods with the highest protein identifications, MagNet, ENRICHplus, EasySep, and EXONET, performed similarly in terms of total number of identified proteins and precursors. All four methods enabled the detection of typical EV proteins while simultaneously enriching cellular proteins and depleting highly abundant soluble plasma proteins. However, despite using plasma samples collected within the same cohort and prepared under identical conditions, we observed variation in the total number of identified proteins and precursors between individual samples. The EV-capturing bead methods (EasySep and EXONET) showed the highest between-sample variability, whereas ENRICHplus exhibited the most consistent performance. Regardless of the enrichment method, all four top-performing enrichment methods increased protein identifications up to 7-fold compared to neat plasma. A key limitation of all tested techniques became evident when we analyzed three commercial plasma samples. Two of these samples yielded poor plasma protein enrichment, whereas a third sample from the same vendor performed comparably to the cohort plasma samples.

MagNet is reported to enrich EV particles by binding negatively charged phospholipids such as phosphatidylserine.^15^ In contrast, EasySep and EXONET rely on antibody-based EV capture without additional physicochemical enrichment. We confirmed that all three approaches enriched typical EV protein markers from the Top 100 EV dataset^40^, many of which are undetectable in neat plasma. ENRICHplus similarly enriched proteins associated with EV particles at intensities comparable to MagNet and EV-capturing beads. The overall protein identification correlation between these four methods across six cohort samples ranged from 0.71 to 0.96, with intra-method correlation mostly between 0.9 and 0.98. Importantly, 80% of all proteins identified in this study were shared among these methods, indicating a largely overlapping protein set, except for ENRICHiST that enriched the lowest number of proteins.

A major challenge in plasma proteomics using enrichment techniques is the influence of pre-enrichment variables, particularly blood collection, on protein identifications. Strikingly, two out of three commercial plasma samples from the same vendor had two-to four-fold lower protein identifications than the plasma samples from our cohort. According to vendor’s protocol (personal communication), these commercial plasma samples were centrifuged at 2900 g for 10 min, while our cohort samples underwent 15 min centrifugation at 1500 g, in accordance with the Early Detection Research Network recommendations^30^ and the MagNet protocol provided by the SAX beads manufacturer, MagReSyn. The number of proteins identified using MagNet in this study (∼4200 proteins) was similar to the previously reported preprint findings,^15^ with 87% of proteins overlapping between datasets. However, a recent study using the Orbitrap Astral MS and plasma centrifuged at 3000 g for 5 min reported only 1200–2200 proteins with MagNet and ENRICHplus, despite employing a chromatographic gradient of similar length to our study.^18^ While we demonstrated that optimizing the plasma-to-bead ratio from 4:1 (as used in previous studies^15,18^) to 40:1 doubled the number of protein identifications, we propose that variations in centrifugation protocols may also play a key role in the differences observed in protein identifications between studies.

Another challenge with the plasma protein enrichment methods is validating the biological significance of the enriched proteins. When comparing our protein identification results to the Human Protein Atlas (HPA) transcriptomics dataset^43^, we found that the most tissue-enriched proteins were predominantly expressed in plasma cells/bone marrow and hepatocytes/liver. This finding aligns with their roles as major sources of soluble blood proteins. Further comparison with two tissue proteomics datasets^44,45^ revealed a subset of proteins highly enriched in platelets. Analysis of ∼2400 proteins shared between the four most effective enrichment methods in this study and the platelet proteome from the Human Proteome Map^45^ showed a moderate correlation between protein intensities. Notably, single-centrifuged plasma samples still contain a substantial number of residual platelets, which can be reduced ten-fold with an additional centrifugation step or fully depleted via high-speed centrifugation.^46,47^ Residual platelets may fragment upon plasma thawing and subsequently become enriched by plasma enrichment methods. Additionally, single-spun plasma releases a high number of CD9- and CD63-positive EVs after thawing, whereas this effect is less pronounced or absent in double-spun plasma.^47,48^ We strongly emphasize the importance of investigating pre-enrichment variables in plasma handling, particularly the number of centrifugation steps and centrifugation speed, as they may directly impact protein identifications with different plasma enrichment methods.

The EV-capturing bead workflows utilized here, EasySep and EXONET, included a high-speed centrifugation step for thawed plasma prior to enrichment. Since this step should deplete any remaining cells or cell fragments, and the EV-capturing enrichment kits resulted in similar total protein identifications in comparison to MagNet and ENRICHplus, it is likely that the platelet-associated proteins detected here may partly originate from EVs released by platelets before blood collection or after plasma freezing and thawing. For neat plasma proteomics studies, and particularly those involving EVs, it has been recommended to use double-spin plasma to deplete platelets.^24,49^ However, even double-spun plasma still retains a significant number of platelets.^50^ Moreover, a high proportion of EVs in blood circulation naturally originate from platelets, megakaryocytes, erythrocytes or immune cells.^51–54^ Therefore, further research is needed to assess how effectively plasma enrichment techniques detect tissue leakage proteins and tissue-derived EVs across different diseases, thereby advancing biomarker research and disease diagnostics. Finally, we acknowledge that biobanks, whether large national collections or smaller clinical trial repositories, contain highly valuable plasma samples, where controlling pre-processing variables is rarely feasible.

The rapid development of new methods to enhance plasma proteome coverage has led to the introduction of many commercial products and academically developed techniques.^4,11–19,23,25^ These methods vary in cost and number of protein identifications. For example, in this study, the list price per sample ranged from 2 dollars for MagNet to 170 dollars for ENRICHplus (Table S7). In addition, advancements in mass spectrometry instrumentation, such as the Orbitrap Astral, will further improve the protein identifications with the plasma enrichment methods.^16,55^ These faster mass spectrometry instruments with shorter gradients may allow sample throughputs of up to 200 SPD.^55^

Increased throughput in LC-MS analysis also demands higher throughput for sample preparation. Here, we developed an automated version of MagNet in a 96-well plate format for the Biomek i5 liquid handler. The workflow fully integrates all steps, starting from plasma sample to LC-MS ready samples loaded in the disposable Evotip trap columns. A similar workflow has previously been reported for the magnetic bead protein aggregation capture workflow integrated with Evotip loading in the Opentrons OT-2 liquid handling robot.^6^ Our MagNet method demonstrated excellent reproducibility, with an average CV of 18% for protein quantification across two full 96-well plates and six different plasma samples. This result is highly comparable to what has been previously reported for the manual MagNet workflow (13% and 21%)^15,18^ or for a semi-automated acid precipitation method for neat plasma (16%)^8^, and better than reported for an automated neat plasma workflow (33%).^29^ The development of the automated MagNet workflow relied on the high binding capacity and rapid magnetization-based recovery of SAX beads, allowing us to miniaturize the bead volume per well to 0.5 µl while simultaneously reducing the amount of digestion enzyme to 90 ng per well. Even after miniaturizing the bead volume, we had to dilute the samples 10-fold before loading them into Evotips to achieve suitable input for our mass spectrometer. Another factor for further optimization is the digestion step to reduce the time required for sample preparation. Most of the time in the current workflow is spent on the overnight digestion at 37 °C. It has been shown that on-bead digestion can be conducted efficiently even within four hours at 37 °C.^6^ Finally, the current automated method could be readily scaled up to process six plates in a single run, depending on the capacity of the liquid handler.

In conclusion, we compared five different plasma protein enrichment methods to neat plasma. The four best-performing enrichment methods (MagNet, ENRICHplus, EasySep and EXONET) increased protein identification 5- to 7-fold in comparison to neat plasma. The highest number of proteins identified from a single plasma sample exceeded 4000 proteins when using a 30 SPD chromatographic method. We further developed a fully automated plasma enrichment method for 20 µl of plasma using the MagNet method, resulting in a remarkable low total sample cost of less than 4$ and excellent reproducibility, with a coefficient of variation below 20% for protein quantification.

## Acknowledgements

Dorte Bekker-Jensen and Magnus Huusfeldt from Evosep Biosystems and Ian Shoemaker from Beckman Coulter are acknowledged for their support in developing the automated Biomek workflows. We thank for the support from Research Council of Finland, Institute of Biotechnology, HiLIFE and Biocenter Finland. E.J. acknowledges funding from the Emil Aaltonen Foundation.

