## Supplementary for "Automated MagNet Enrichment Unlocks Deep and Cost-Effective LC-MS Plasma Proteomics"

**CONTENTS**

**TABLES**

Table S1. Biomek i5 workflow for the automated MagNet protocol.

Table S2. MS and TIMS settings for all methods.

Table S3. MS/MS settings for 60 SPD and 100 SPD methods

Table S4. MS/MS settings for the 30 SPD method.

Table S5. dia-PASEF windows for 60 SPD and 100 SPD methods.

Table S6. dia-PASEF windows for the 30 SPD method.

Table S7. Price comparison of plasma enrichment methods.

**FIGURES**

Figure S1. Total ion chromatograms (TIC) of a sample from all five enrichment methods and neat plasma.

Figure S2. Plasma enrichment methods with commercial samples included.

Figure S3. Comparison of plasma enrichment methods and neat plasma to PeptideAtlas.

Figure S4. Comparison of proteins detected in this study with previous publications.

Figure S5. Comparison of proteins detected in this study with previous plasma EV proteomics publications.

Figure S6. Venn diagrams showing overlapping proteins between plasma enrichment methods or neat plasma and two commercial targeted protein assays: Olink HT and SomaScan 10k.

Figure S7. Gene ontology (GO) analysis of proteins in four clusters.

Figure S8. Pathway and protein class analysis for proteins in four clusters.

Figure S9. Cell-type enriched proteins based on The Human Protein Atlas (HPA) single-cell transcriptomics dataset.

Figure S10. Tissue expression of proteins identified in study.

Figure S11. Tissue expression of proteins identified in this study using The Human Proteome Map (HPM).

Figure S12. Automated Evotip loading and MagNet workflow using a Biomek liquid handler.

**Table S1. Biomek i5 workflow for the automated MagNet protocol.** All pipetting steps utilize 90 µl non-sterile tips from Beckman Coulter (product number B85881). For all steps, except sample dilution, a same tip box was used for pipetting buffers and solvents, and another tip box for removing liquid from sample plate.

| **PLASMA BINDING** |
|---|
| 1. Shake the sample plate for 1 min at 1800 RPM.<br>2. Dispense 50 µl of wash buffer to the wells of sample plate.<br>3. Shake the sample plate for 1 min at 1650 RPM.<br>4. Transfer the plate onto magnet plate and completely remove liquid from the wells. |
| **BEAD WASHING (repeat 3 times)** |
| 1. Dispense 120 µl of wash buffer to the well of sample plate.<br>2. Shake the sample plate for 1 min a 1650 RPM.<br>3. Transfer the plate onto magnet plate and completely remove liquid from the wells. |
| **LYSIS, REDUCTION and ALKYLATION** |
| 1. Aspirate 50 µl of wash buffer (for rinsing the tips) followed by a 10 µl air gap and 15 µl of lysis buffer. Transfer 15 µl of lysis buffer to the wells of sample plate and dispense the rest of liquid to waste.<br>2. Shake the sample plate for 30 s at 1800 RPM.<br>3. Incubate the sample plate for 1 h at 37 °C under 1600 RPM shaking. |
| **PROTEIN AGGREGATION CAPTURE** |
| 1. Dispense 40 µl of isopropanol to the wells of sample plate.<br>2. Shake the sample plate for 30 s at 1700 RPM.<br>3. Pause the protocol for 10 min.<br>4. Shake the sample plate for 30 s at 1700 RPM.<br>5. Transfer the plate onto magnet plate and completely remove liquid from the wells. |
| **BEAD WASHING** |
| 1. Dispense 150 µl of isopropanol to the wells of sample plate.<br>2. Transfer the plate onto magnet plate and completely remove liquid from the wells.<br>3. Dispense 75 µl of isopropanol to the wells of sample plate.<br>4. Shake the sample plate for 30 s at 1500 RPM.<br>5. Transfer the plate onto magnet plate and completely remove liquid from the wells.<br>6. Dispense 180 µl of isopropanol to the wells of sample plate.<br>7. Transfer the plate onto magnet plate and completely remove liquid from the wells. |

| DIGESTION |
| --- |
| 1. Wash the tips twice by aspirating 90 µl of wash buffer and dispensing it to waste.<br>2. Aspirate 50 µl of digestion buffer and dispense it to the wells of sample plate.<br>3. Shake the sample plate for 1 min at 1800 RPM.<br>4. Seal the plate tightly with a sealing foil (product number AB0626 from Thermo Fisher Scientific).<br>5. Incubate the sample plate for 16 h at 37 °C under 1100 RPM shaking.<br>6. After incubation, remove the sealing foil from the sample plate |
| **SAMPLE DILUTION** |
| 1. Transfer 90 µl of Solvent A with new tips to a new LoBind 96-well PCR plate.<br>2. Shake the sample plate for 2 min at 1600 RPM.<br>3. Transfer 10 µl from the sample plate with new tips to the dilution plate, and tip-mix the sample five times with 20 µl volume after dispensing it in the dilution plate.<br>4. Shake the dilution plate for 2 min at 1700 RPM.<br>5. Proceed to Evotip loading. |

**Table S2. MS and TIMS settings for all methods.**

| | |
|---|---|
| Scan Begin | 100 m/z |
| Scan End | 1 700 m/z |
| Ion Polarity | Positive |
| Scan Mode | dia-PASEF |
| Ramp Time | 100 ms |
| Accumulation Time | 100 ms |
| Duty Cycle | 100% |
| Ramp Rate | 9.42 Hz |

**Table S3. MS/MS settings for the 60 SPD and 100 SPD methods**

| | |
|---|---|
| Cycle Time Estimate | 1.06 s |
| Number of MS1 Ramps | 1 |
| Number of MS/MS Ramps | 9 |
| Number of MS/MS Windows | 21 |
| Mass Range | 475.0 to 1000.0 Da |
| Mobility Range | 0.85 to 1.27 1/K0 |
| Collision Energy Settings | 20 eV (0.60 1/K0) to 59.0 eV (1.60 1/K0) |

**Table S4. MS/MS settings for the 30 SPD method.**

| | |
|---|---|
| Cycle Time Estimate | 1.80 s |
| Number of MS1 Ramps | 1 |
| Number of MS/MS Ramps | 16 |
| Number of MS/MS Windows | 42 |
| Mass Range | 294.5 to 1324.0 Da |
| Mobility Range | 0.73 to 1.33 1/K0 |
| Collision Energy Settings | 20 eV (0.60 1/K0) to 59.0 eV (1.60 1/K0) |

**Table S5. dia-PASEF windows for the 60 SPD and 100 SPD methods.**

| #MS Type | Cycle Id | Start IM [1/K0] | End IM [1/K0] | Start Mass [m/z] | End Mass [m/z] |
|---|---|---|---|---|---|
| MS1 | 0 | - | - | - | - |
| PASEF | 1 | 0.85 | 0.8911 | 475 | 500 |
| PASEF | 1 | 0.9088 | 1.0604 | 695.5 | 720.5 |
| PASEF | 1 | 1.078 | 1.2296 | 916 | 941 |
| PASEF | 2 | 0.85 | 0.9099 | 499.5 | 524.5 |
| PASEF | 2 | 0.9276 | 1.0792 | 720 | 745 |
| PASEF | 2 | 1.0968 | 1.2484 | 940.5 | 965.5 |
| PASEF | 3 | 0.85 | 0.9287 | 524 | 549 |
| PASEF | 3 | 0.9464 | 1.098 | 744.5 | 769.5 |
| PASEF | 3 | 1.1156 | 1.2672 | 965 | 1000 |
| PASEF | 4 | 0.85 | 0.9475 | 548.5 | 573.5 |
| PASEF | 4 | 0.9652 | 1.1168 | 769 | 794 |
| PASEF | 5 | 0.85 | 0.9664 | 573 | 598 |
| PASEF | 5 | 0.984 | 1.1356 | 793.5 | 818.5 |
| PASEF | 6 | 0.85 | 0.9852 | 597.5 | 622.5 |
| PASEF | 6 | 1.0028 | 1.1544 | 818 | 843 |
| PASEF | 7 | 0.85 | 1.004 | 622 | 647 |
| PASEF | 7 | 1.0216 | 1.1732 | 842.5 | 867.5 |
| PASEF | 8 | 0.87 | 1.0228 | 646.5 | 671.5 |
| PASEF | 8 | 1.0404 | 1.192 | 867 | 892 |
| PASEF | 9 | 0.89 | 1.0416 | 671 | 696 |
| PASEF | 9 | 1.0592 | 1.2108 | 891.5 | 916.5 |

**Table S6. dia-PASEF windows for the 30 SPD method.**

| #MS Type | Cycle Id | Start IM [1/K0] | End IM [1/K0] | Start Mass [m/z] | End Mass [m/z] |
|---|---|---|---|---|---|
| MS1 | 0 | - | - | - | - |
| PASEF | 1 | 0.73 | 0.7707 | 294.5 | 319.5 |
| PASEF | 1 | 0.7888 | 1.0929 | 686.5 | 711.5 |
| PASEF | 1 | 1.111 | 1.33 | 1078.5 | 1103.5 |
| PASEF | 2 | 0.73 | 0.7908 | 319 | 344 |
| PASEF | 2 | 0.8089 | 1.113 | 711 | 736 |
| PASEF | 2 | 1.1311 | 1.33 | 1103 | 1128 |
| PASEF | 3 | 0.73 | 0.811 | 343.5 | 368.5 |
| PASEF | 3 | 0.829 | 1.1332 | 735.5 | 760.5 |
| PASEF | 3 | 1.1512 | 1.33 | 1127.5 | 1152.5 |
| PASEF | 4 | 0.73 | 0.8311 | 368 | 393 |
| PASEF | 4 | 0.8492 | 1.1533 | 760 | 785 |
| PASEF | 4 | 1.1714 | 1.33 | 1152 | 1177 |
| PASEF | 5 | 0.73 | 0.8512 | 392.5 | 417.5 |
| PASEF | 5 | 0.8693 | 1.1734 | 784.5 | 809.5 |
| PASEF | 5 | 1.1915 | 1.33 | 1176.5 | 1201.5 |
| PASEF | 6 | 0.73 | 0.8714 | 417 | 442 |
| PASEF | 6 | 0.8895 | 1.1936 | 809 | 834 |
| PASEF | 6 | 1.2116 | 1.33 | 1201 | 1226 |
| PASEF | 7 | 0.73 | 0.8915 | 441.5 | 466.5 |
| PASEF | 7 | 0.9096 | 1.2137 | 833.5 | 858.5 |
| PASEF | 7 | 1.2318 | 1.33 | 1225.5 | 1250.5 |
| PASEF | 8 | 0.73 | 0.9116 | 466 | 491 |
| PASEF | 8 | 0.9297 | 1.2338 | 858 | 883 |
| PASEF | 8 | 1.2519 | 1.33 | 1250 | 1275 |
| PASEF | 9 | 0.73 | 0.9318 | 490.5 | 515.5 |
| PASEF | 9 | 0.9499 | 1.254 | 882.5 | 907.5 |
| PASEF | 9 | 1.2721 | 1.33 | 1274.5 | 1299.5 |
| PASEF | 10 | 0.73 | 0.9519 | 515 | 540 |
| PASEF | 10 | 0.97 | 1.2741 | 907 | 932 |
| PASEF | 10 | 1.2922 | 1.33 | 1299 | 1324 |
| PASEF | 11 | 0.73 | 0.9721 | 539.5 | 564.5 |
| PASEF | 11 | 0.9901 | 1.2942 | 931.5 | 956.5 |
| PASEF | 12 | 0.73 | 0.9922 | 564 | 589 |
| PASEF | 12 | 1.0103 | 1.3144 | 956 | 981 |
| PASEF | 13 | 0.73 | 1.0123 | 588.5 | 613.5 |
| PASEF | 13 | 1.0304 | 1.33 | 980.5 | 1005.5 |
| PASEF | 14 | 0.73 | 1.0325 | 613 | 638 |
| PASEF | 14 | 1.0505 | 1.33 | 1005 | 1030 |
| PASEF | 15 | 0.7485 | 1.0526 | 637.5 | 662.5 |
| PASEF | 15 | 1.0707 | 1.33 | 1029.5 | 1054.5 |
| PASEF | 16 | 0.7686 | 1.0727 | 662 | 687 |
| PASEF | 16 | 1.0908 | 1.33 | 1054 | 1079 |

**Table S7. Price comparison of plasma enrichment methods.** The prices are calculated based on the list prices of highest package sizes for all reagents, no taxes or shipping costs are included.

| **Enrichment method** | **Cost per sample** |
|---|---|
| ENRICHplus | 169$ (kit) |
| ENRICHiST | 93$ (kit) |
| EasySep | 38$ (kit, S-Trap purification column and digestion enzyme) |
| EXONET | 59$ (kit, S-Trap purification column and digestion enzyme) |
| MagNet, manual | 1.9$ (beads and digestion enzyme) |
| MagNet, automated | 0.28$ (beads and digestion enzyme), or total cost of 3.6$ (including also Biomek tips, 96 well plate and Evotips) |

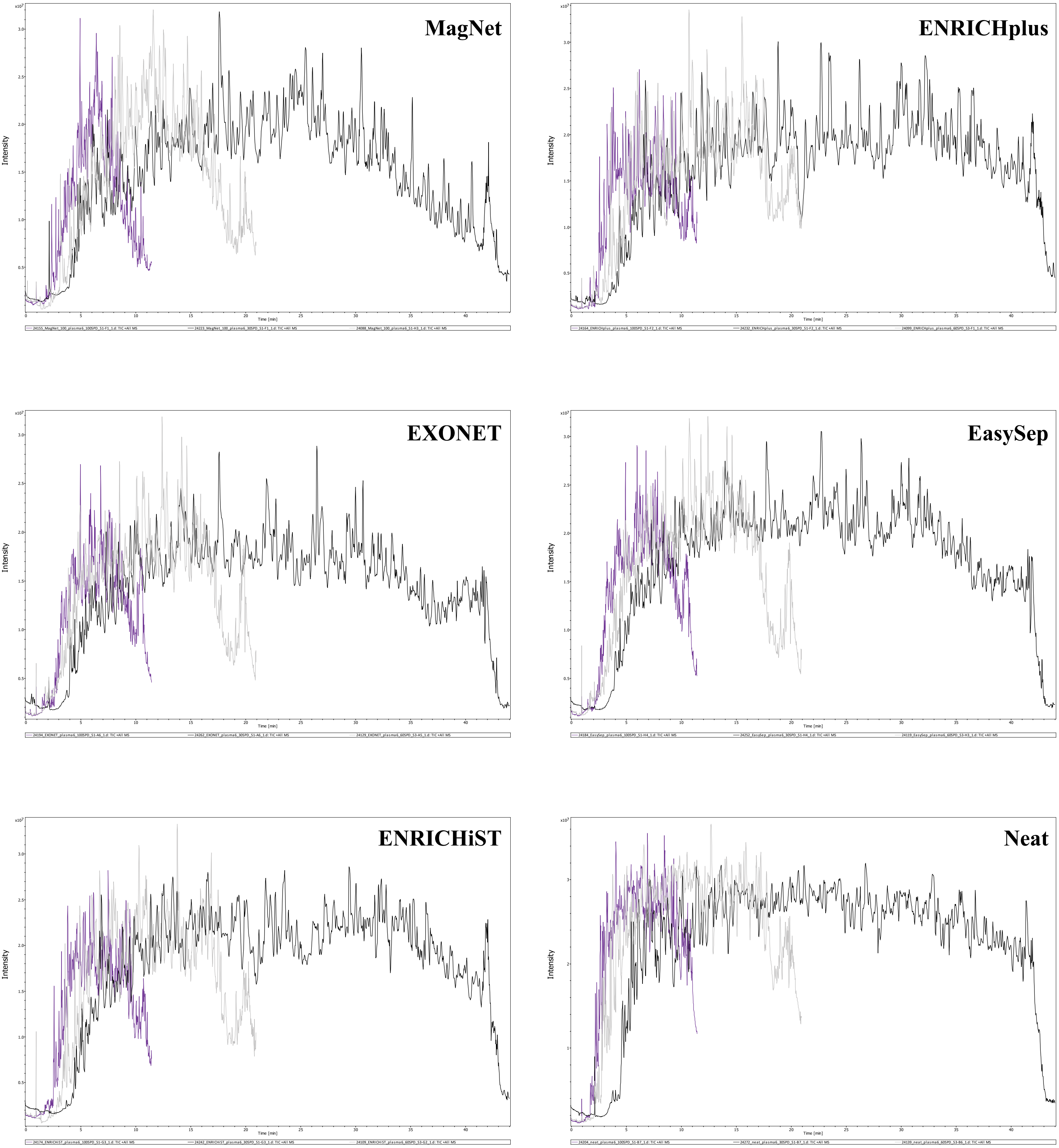


**Figure S1**. **Total ion chromatograms (TIC) of a sample from all five enrichment methods and neat plasma.** Black, grey, and purple lines represent TICs corresponding to the 30, 60, and 100 SPD enrichment methods, respectively.

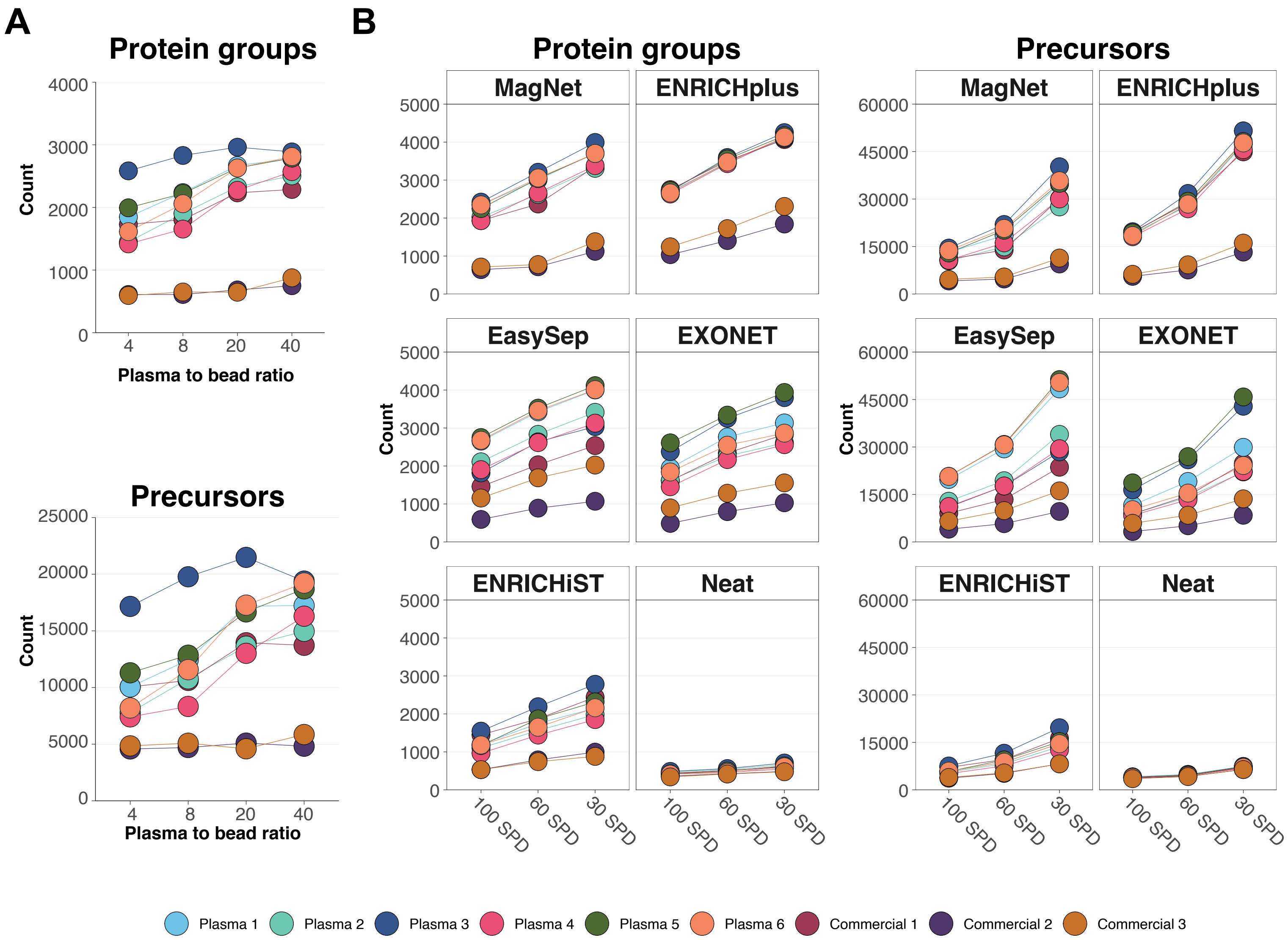


**Figure S2**. **Plasma enrichment methods with commercial samples included.** The same results shown in Figure 1 are presented here, with the addition of three different commercial plasma samples.
(**A**) Optimization of the plasma-to-bead ratio for MagNet using the 60 SPD chromatographic gradient method. The 40:1 plasma-to-bead volume ratio was selected for MagNet samples presented in panel B.
(**B**) Five different enrichment methods and the neat plasma workflow were applied to six different plasma samples, which were analyzed using three different chromatographic gradients.

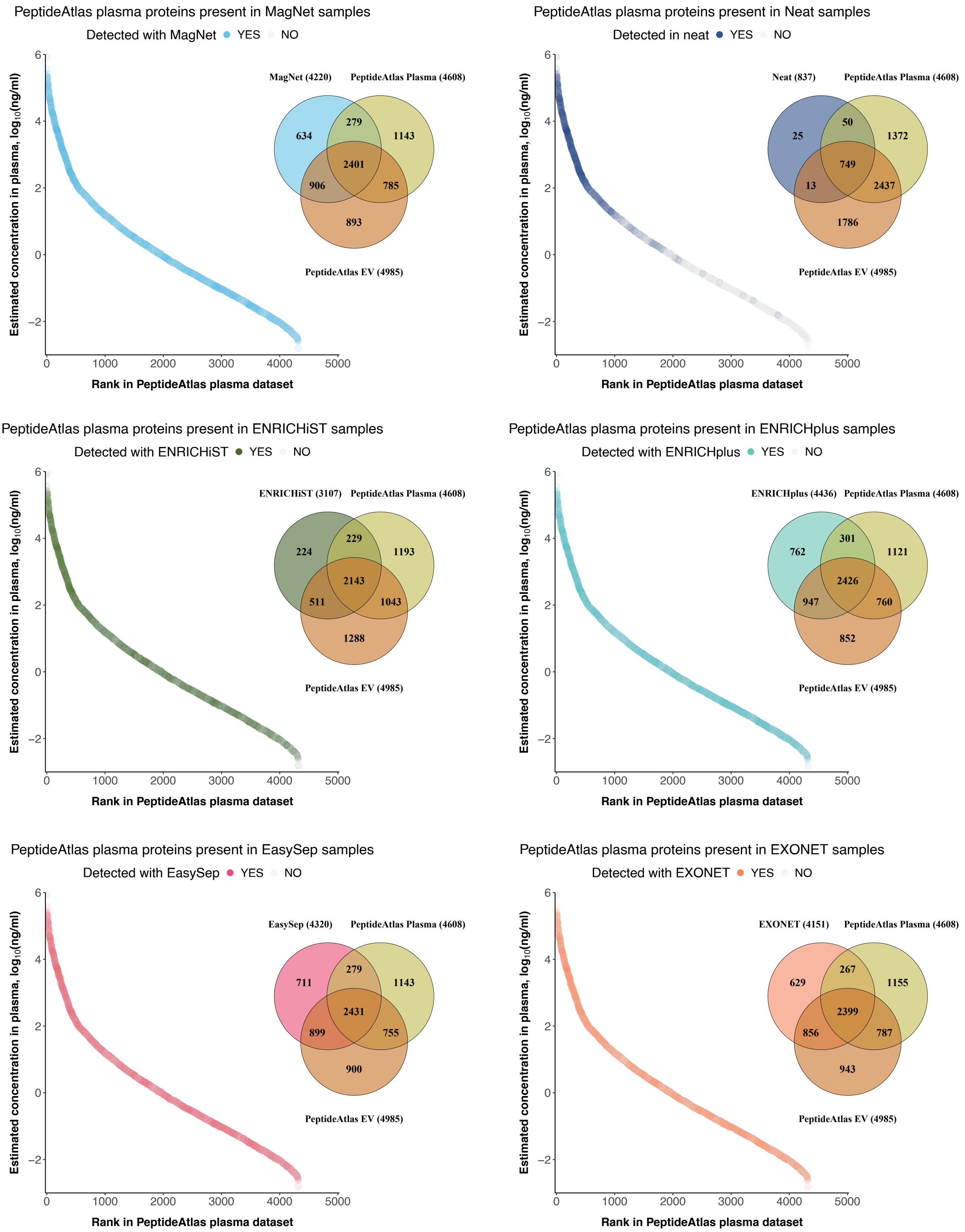


**Figure S3**. **Comparison of plasma enrichment methods and neat plasma to PeptideAtlas.** Proteins identified by each method were compared to their rankings and estimated plasma concentrations (ng/mL) in the PeptideAtlas Human Plasma 2023-04 Build. Colored dots indicate proteins detected in both datasets, while grey dots represent proteins found only in PeptideAtlas. Venn diagrams show the total number of overlapping proteins between each enrichment method or neat plasma, and the PeptideAtlas Human Plasma and Human Plasma Extracellular Vesicle (EV) datasets. The total number of proteins in each dataset is shown in parentheses. PeptideAtlas builds (Human Plasma 2023-04 and Human Plasma Extracellular Vesicle 2023-04) were downloaded from the PeptideAtlas (https://peptideatlas.org/).

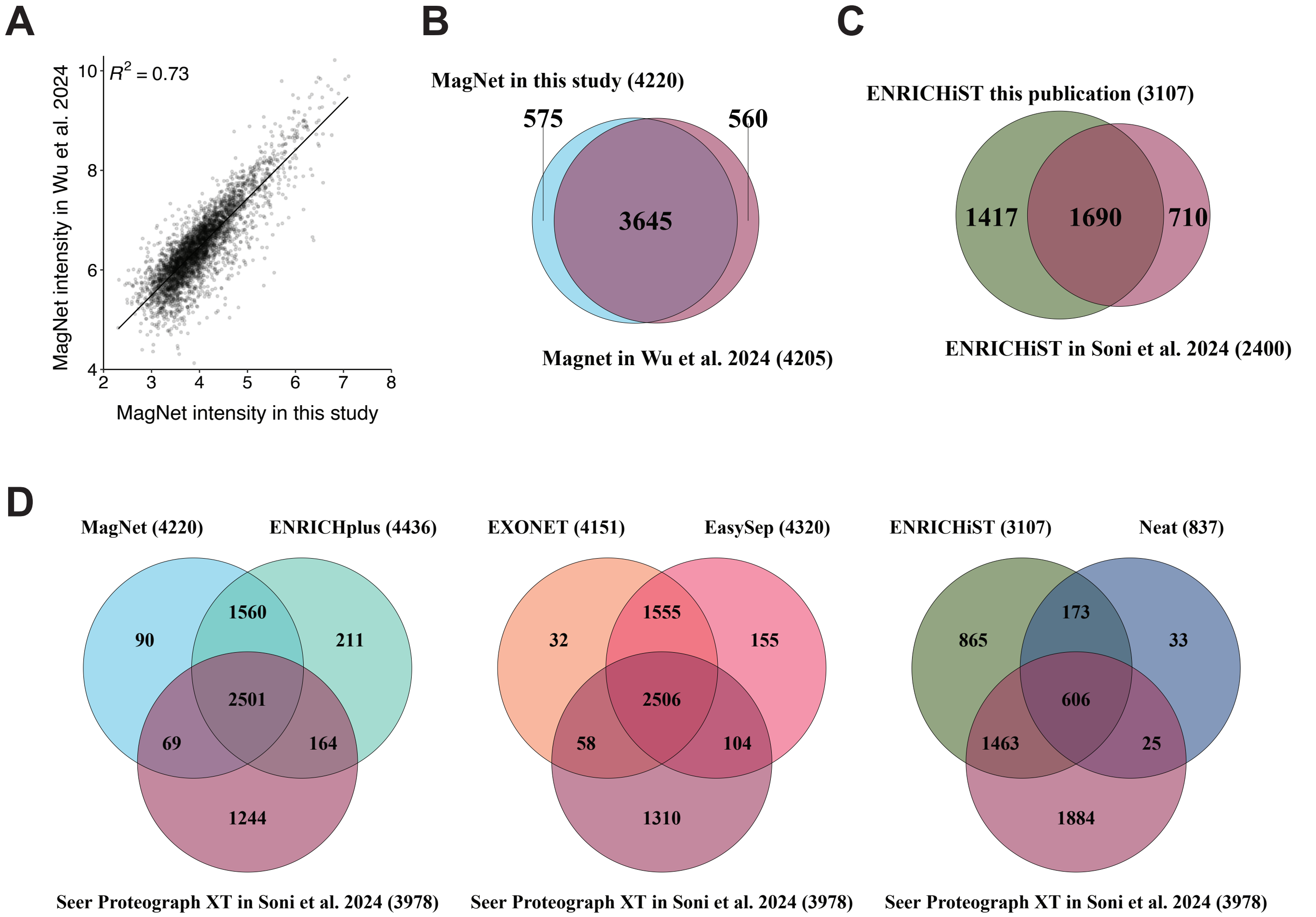


**Figure S4**. **Comparison of proteins detected in this study with previous publications. (A)** Linear correlation of $log_{10}$-transformed protein intensities between this study and Wu et al. 2024 (https://doi.org/10.1101/2023.06.10.544439) using the MagNet enrichment method. **(B)** Venn diagram showing overlapping protein identifications between MagNet-based studies. **(C)** Venn diagram of proteins identified with ENRICHiST in this study and in Soni et al. 2024 (https://doi.org/10.1186/s12014-024-09497-2). **(D)** Venn diagrams comparing proteins identified with the enrichment methods in this study to those detected using the Seer Proteograph XT method in Soni et al. 2024. The total protein counts for each dataset are shown in parentheses.

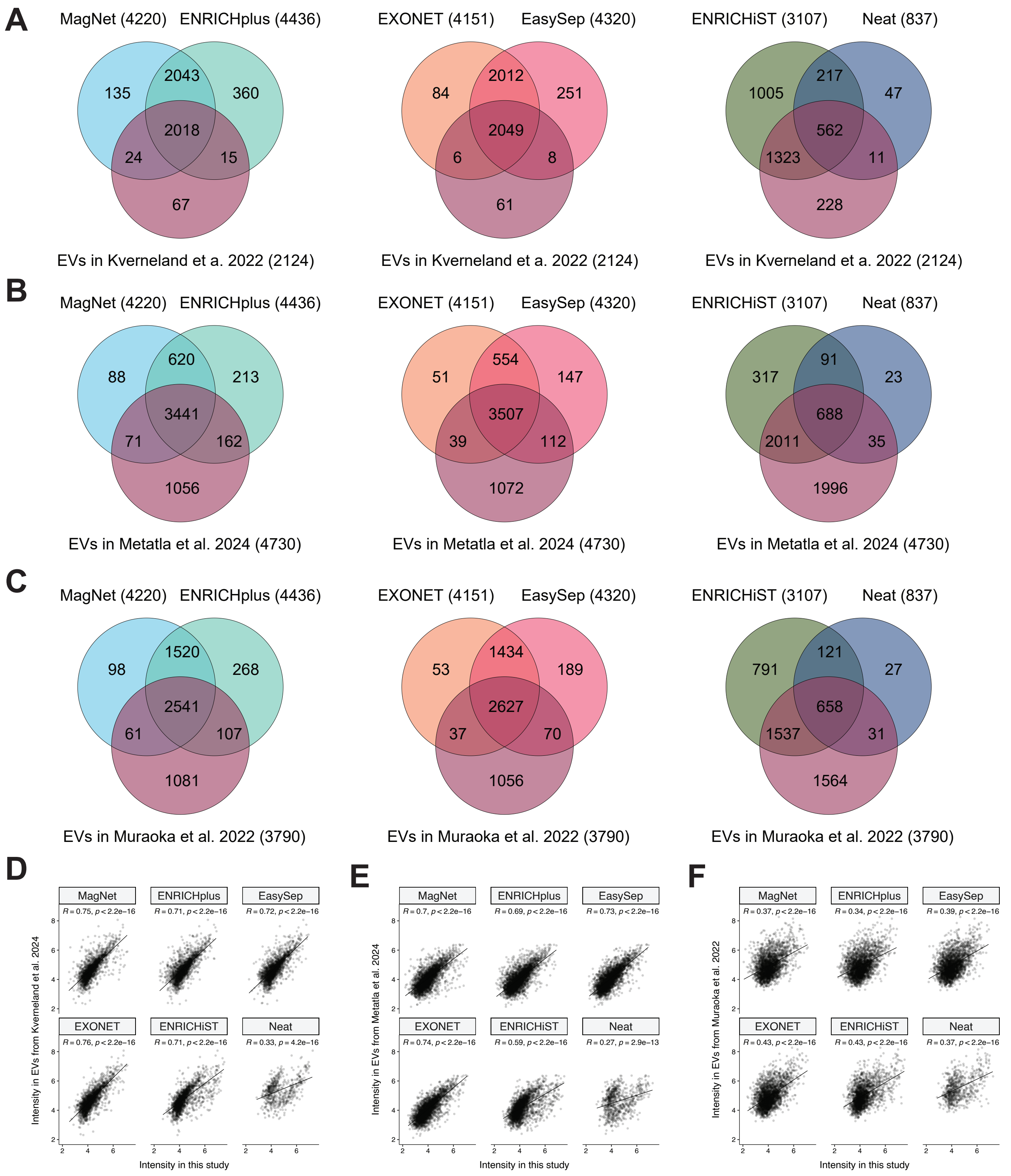


**Figure S5**. **Comparison of proteins detected in this study with previous plasma EV proteomics publications. (A–C)** Venn diagrams show the overlap of proteins identified in this study and three previous studies analyzing plasma EVs. **(A, D)** Comparison with EV proteins identified by Kverneland et al. 2022 (https://doi.org/10.1002/pmic.202200039), using platelet-poor plasma prepared by centrifugation at 20,000 × g. **(B, E)** Comparison with EV proteins identified by Metatla et al. 2024 (https://doi.org/10.1186/s12014-024-09477-6), using platelet-poor plasma prepared by 20,000 × g centrifugation. **(C, F)** Comparison with EV proteins identified by Muraoka et al. 2022 (https://doi.org/10.1016/j.isci.2022.104012), using plasma filtered through a 0.45 µm membrane and isolated by TIM4-affinity capture targeting phosphatidylserine. **(D–F)** Linear correlations of $\log_{10}$-transformed intensities for shared proteins between this study and the respective publications shown in panels A–C.

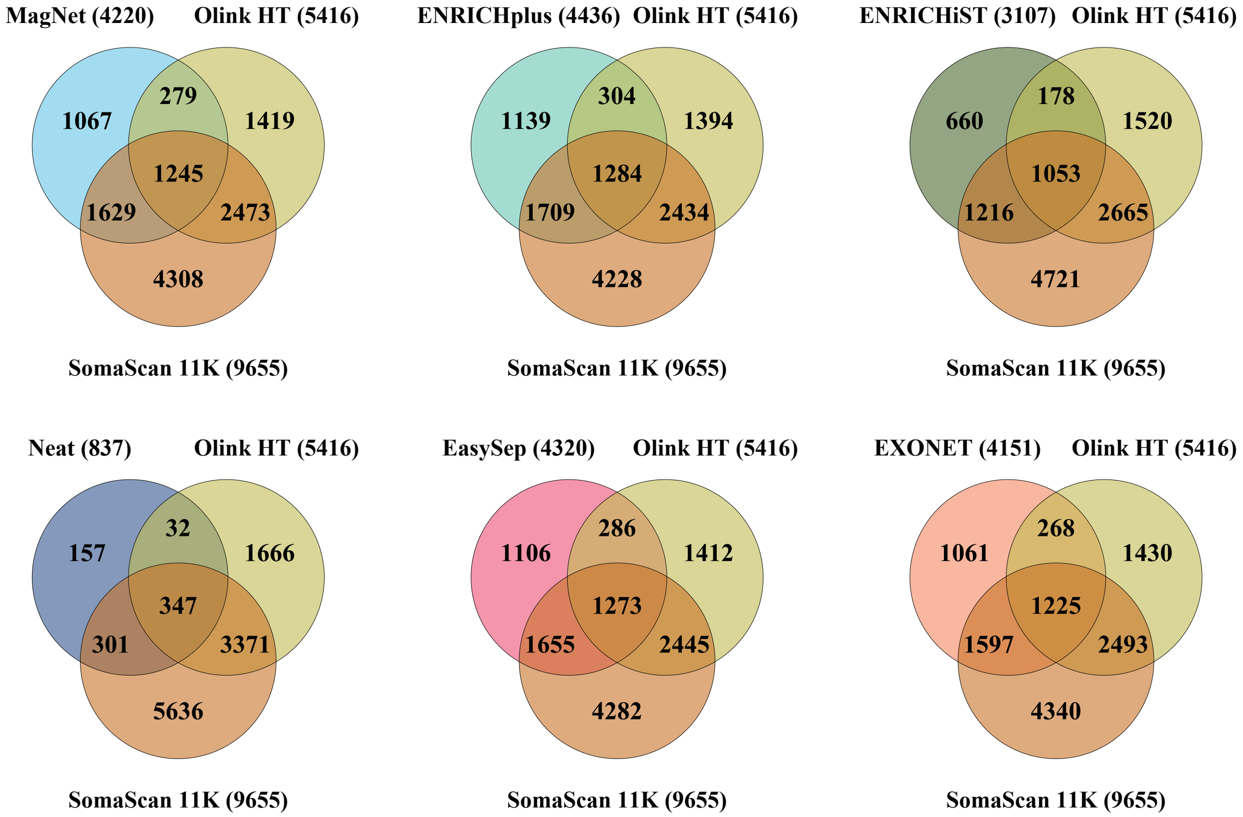


**Figure S6. . Venn diagrams showing overlapping proteins between plasma enrichment methods or neat plasma and two commercial targeted protein assays: Olink HT and SomaScan 10k.** The total number of proteins in each dataset is indicated in parentheses. Protein lists for the targeted assays were obtained from the manufacturers' websites (https://olink.com/ and https://somalogic.com/)

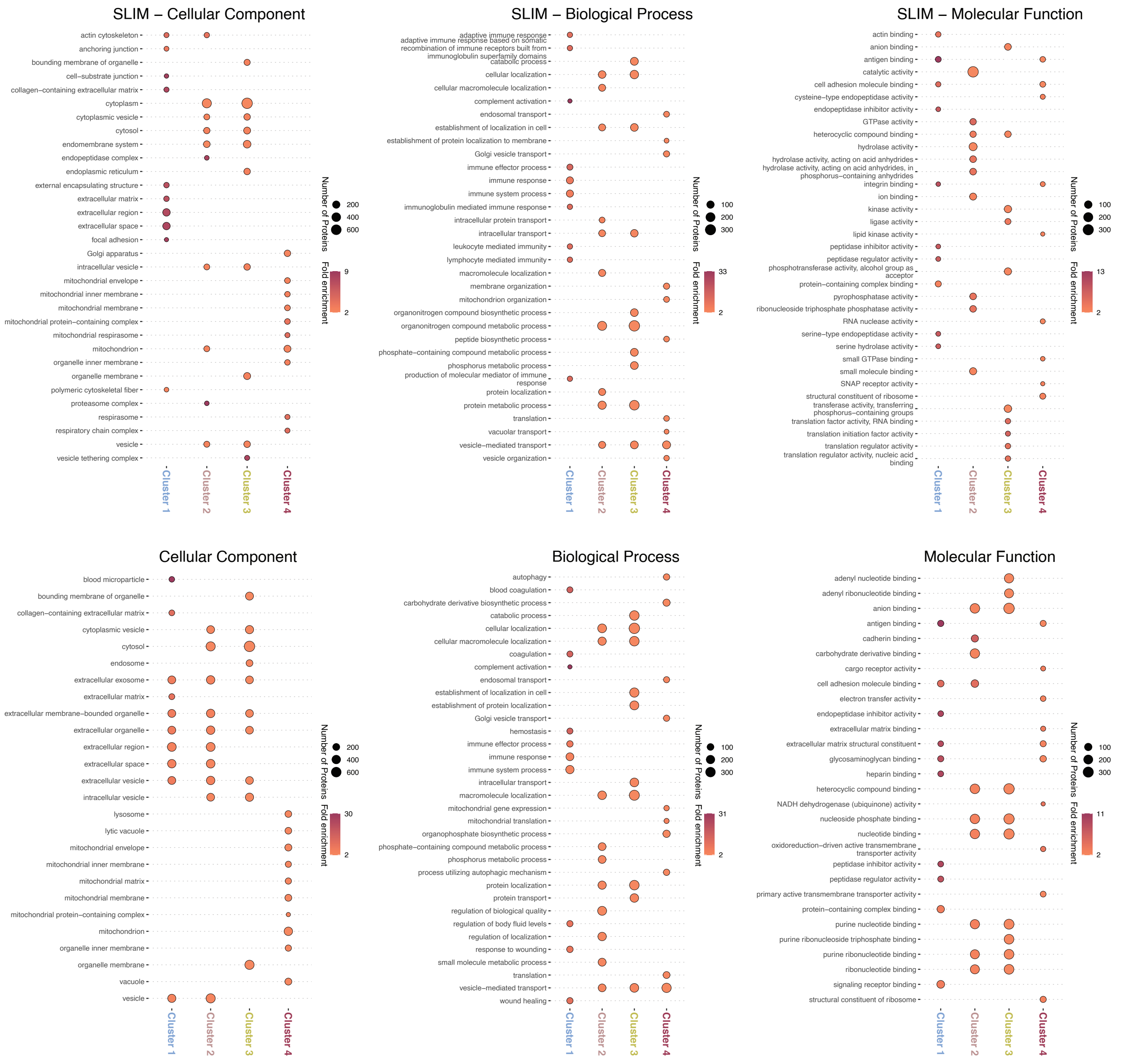


**Figure S7. Gene ontology (GO) analysis of proteins in four clusters. Cluster 1 includes proteins present in neat plasma and shared across all enrichment methods. Cluster 2 contains proteins identified in all five enrichment methods. Cluster 3 comprises proteins with medium intensity, mostly detected in all methods except ENRICHiST and neat plasma. Cluster 4 consists primarily of low-intensity proteins.** Clustering analysis is shown in Figure 3A. GO enrichment analysis was performed using the PANTHER GO-Slim and GO databases (https://pantherdb.org/). Results were filtered to include only terms with greater than two-fold enrichment and at least ten proteins per term. The top ten GO terms with the lowest FDR values were selected for each cluster.

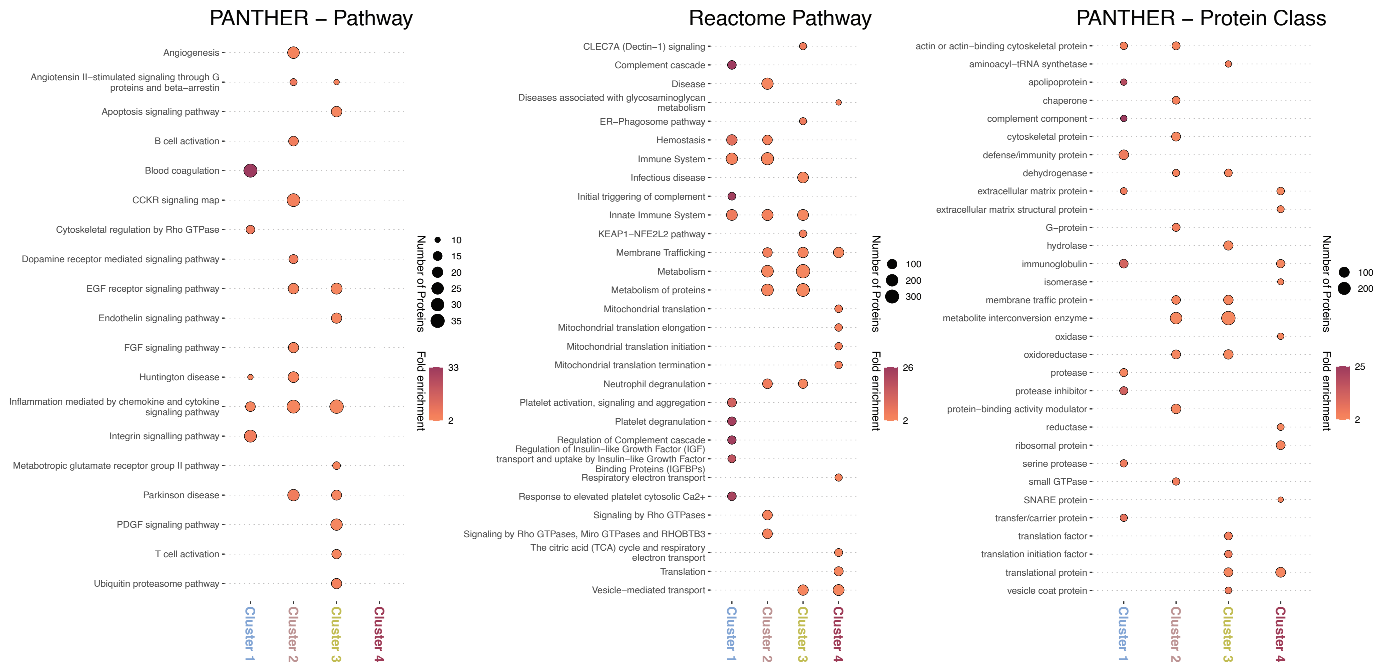


**Figure S8**. **Pathway and protein class analysis for proteins in four clusters. Cluster 1 includes proteins present in neat plasma and shared across all enrichment methods. Cluster 2 contains proteins identified in all five enrichment methods. Cluster 3 comprises proteins with medium intensity, mostly detected in all methods except ENRICHiST and neat plasma. Cluster 4 consists primarily of low-intensity proteins.** Clustering analysis is shown in Figure 3A. Enrichment analysis of proteins in each cluster was performed using PANTHER Pathway, PANTHER Protein Class, and Reactome Pathway annotations (https://pantherdb.org/). Results were filtered to retain terms with greater than two-fold enrichment and at least ten proteins per term. The top ten terms with the lowest FDR values

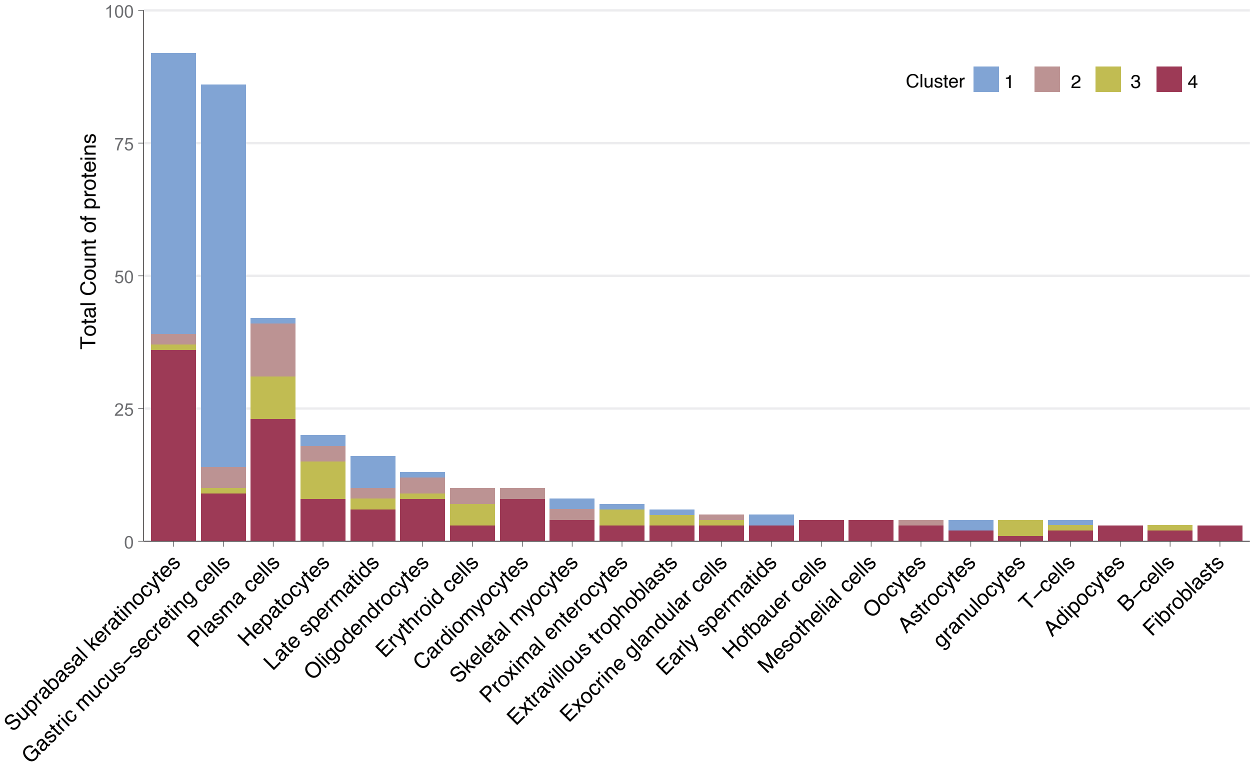


**Figure S9**. **Cell-type enriched proteins based on The Human Protein Atlas (HPA) single-cell transcriptomics dataset.** Clustering analysis is shown in Figure 3A. Only cell types with at least four enriched proteins are displayed. (https://www.proteinatlas.org/)

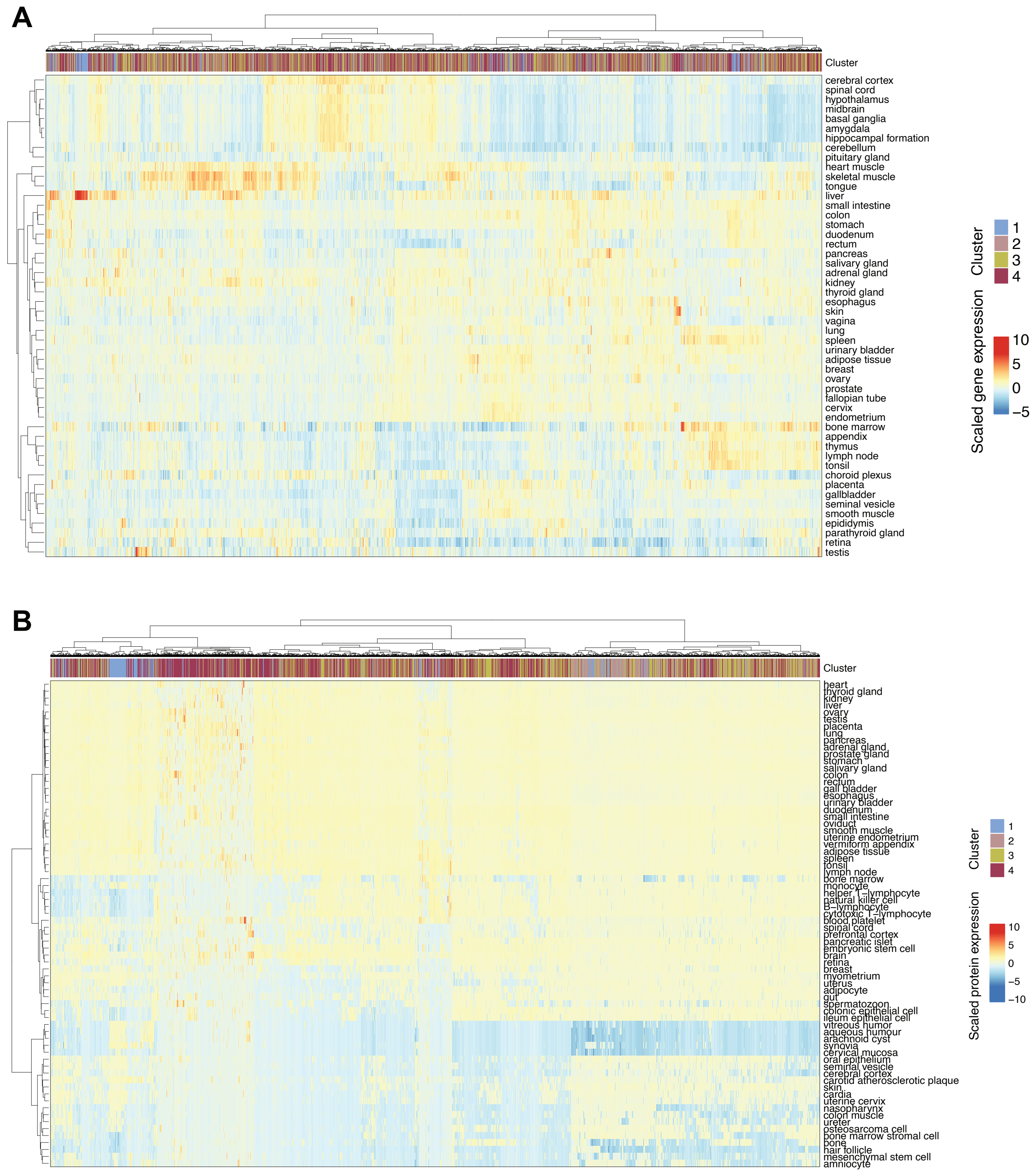


**Figure S10**. **Tissue expression of proteins identified in this study. (A)** Transcript-level expression of the identified proteins was obtained from The Human Protein Atlas (HPA) RNA expression database (https://www.proteinatlas.org/). **(B)** Protein-level expression was obtained from ProteomicsDB (https://www.proteomicsdb.org/). For both panels, expression values were Z-score normalized (mean-centered and scaled by standard deviation) for each gene or protein across tissues. Heatmaps display clustered tissues and genes/proteins. Protein clusters from Figure 3A are indicated in the top panel of each heatmap.

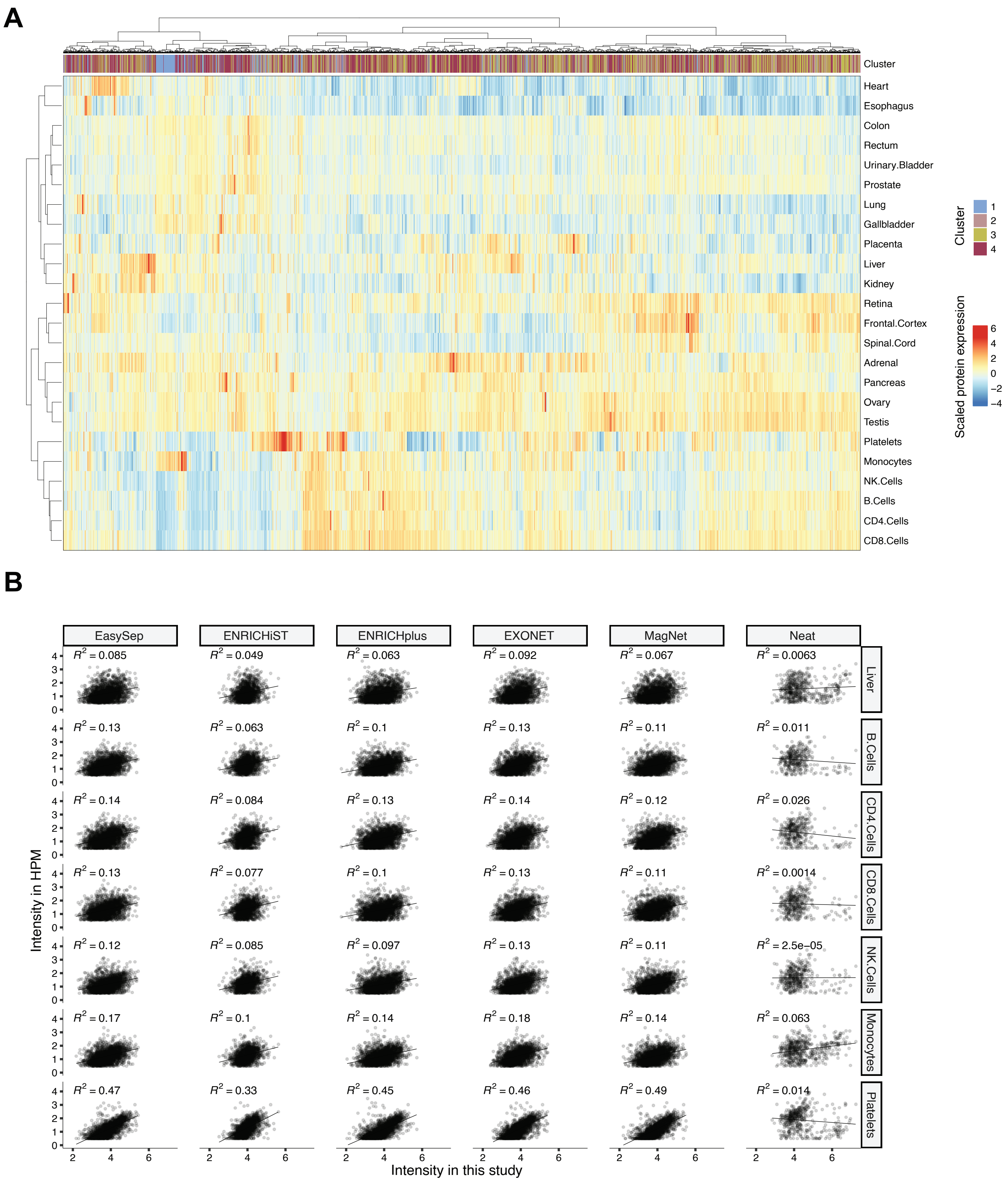


**Figure S11**. **Tissue expression of proteins identified in this study using The Human Proteome Map (HPM).** **(A)** Tissue expression data for the identified proteins were obtained from HPM (http://www.humanproteomemap.org/). Expression values were Z-score normalized (mean-centered and scaled by standard deviation) across tissues. Both tissues and proteins were clustered in the heatmap. Protein clusters from Figure 3A are shown in the top panel. **(B)** Linear correlation of $\log_{10}$-transformed protein intensities between this study and HPM, focusing on blood cell and liver proteomes. Mean intensity values for enriched proteins (those absent from neat plasma) were calculated for each enrichment method, while mean intensities for all proteins in neat plasma were averaged across plasma samples 1–6. Proteins shared between this study and HPM were used for correlation analysis. The average number of shared proteins was 2354 (MagNet), 2477 (ENRICHplus), 2377 (EasySep), 2288 (EXONET), 1657 (ENRICHiST), and 432 (Neat plasma).

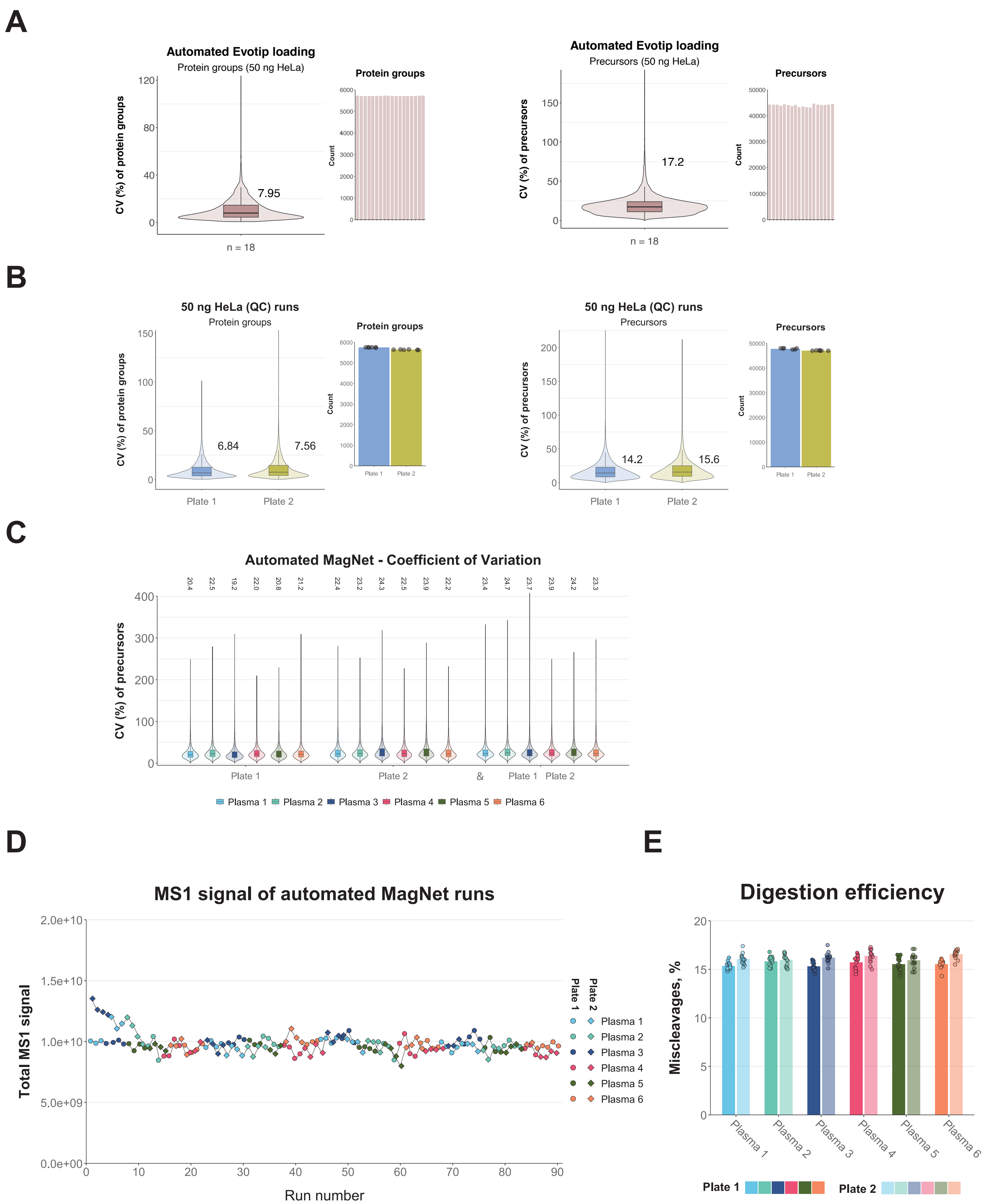


**Figure S12**. **Automated Evotip loading and MagNet workflow using a Biomek liquid handler. (A)** Coefficient of variation (CV, %) for protein groups and precursors in 50 ng HeLa peptide samples (n = 18) processed with automated Evotip loading using a Biomek i5 liquid handler. **(B)** CV (%) of protein groups and precursors in 50 ng HeLa peptide samples processed manually and used as quality control (QC) samples during LC-MS analysis of MagNet sample batches on plate 1 and plate 2. **(C)** CV (%) of precursors within each plasma sample across plate 1 and plate 2 (n = 15 per plate; n = 30 total). **(D)** Total MS1 signal, reported by DIA-NN software, per plasma sample in plate 1 and plate 2. **(E)** Digestion efficiency for all plasma samples in plate 1 and plate 2, measured as the relative abundance of peptides with one missed cleavage.
